# Inhibition of HDAC7 reprograms the histone H3.3 landscape to induce heterochromatin spreading and DNA replication defects in cancer cells

**DOI:** 10.1101/2024.03.12.584656

**Authors:** Ola Hassan, Mattia Pizzagalli, Owen Leary, John P. Zepecki, Adrianne Corseri, Laura Jinxuan Wu, Shiven Sasipalli, Daniel Lee, Lindsey Hayward, Lily Tran, Eduardo Fajardo, Andras Fiser, David Karambizi, Nikos Tapinos

## Abstract

Class IIa histone deacetylases (HDACs) are a family of enzymes with minimal histone deacetylase activity but can function as multi-protein interaction hubs. Here we demonstrate the expression of HDAC7, a Class IIa HDAC family member, in glioblastoma tumor tissue from 84 patients, patient-derived glioma stem cells (GSCs) from six patients, and pediatric diffuse pontine glioma (DIPG) cells from three patients. HDAC7 binds to Histone H3.3 and interacts with H3.3 and HIRA on chromatin. Targeted downregulation of HDAC7 expression with a subtype-specific siRNA inhibits the interaction of H3.3 with HIRA while increasing the association of H3.3 with DAXX and H3K9me3. This results in H3.3 being deposited on H3K9me3+/DAPI+ heterochromatin nuclear foci. Inhibition of HDAC7 triggers H3K9me3+ heterochromatin spreading, increased H3K9me3 binding in the cancer genome, and significant alterations in gene expression. Using single molecule DNA fiber approach, we show that HDAC7 inhibition results in a significant increase in replication fork speed without affecting fork symmetry. This altered replication fork speed leads to replication stress, evidenced by phosphorylation of RPA2 and impact on global DNA synthesis, resulting in reduced EdU incorporation. Finally, HDAC7 depletion leads to reduced BRCA2 expression and increased sensitivity of cancer cells to DNA damaging agents. Taken together, these studies uncover the involvement of HDAC7 in the euchromatic H3.3 chaperone network and the effect of HDAC7 depletion on chromatin dynamics, inducing epigenetic restriction and DNA damage in cancer cells.

## Introduction

Epigenetic mechanisms have emerged as key players in cancer development. Alterations in chromatin mediate cell reprogramming and oncogenesis, while epigenetic changes triggered by the tumor microenvironment regulate cancer cell phenotypic transitions and tumor architecture ^1,2^. Chromatin exists in two higher order states: euchromatin refers to loosely packed chromatin that is easily accessible to transcriptional regulators and RNA polymerase, while heterochromatin is densely packed and not accessible ^3,4^. Alteration of the state of chromatin impacts global gene expression by making genes either more or less accessible for transcription.

Interconversion between these two chromatin states occurs through the function of several regulators that affect levels of DNA methylation or posttranslational modification of histones (acetylation, methylation, phosphorylation, sumoylation etc.). This dynamic regulatory system is commonly referred as the epigenetic control of gene transcription ^5^.

Histone deacetylases (HDACs) remove acetyl- groups from histone molecules hence regulating posttranslational modification of histone marks. Expression of HDACs is elevated in several tumors resulting in transcriptional activation of oncogenes, increased transcriptional rates and chromosomal translocations ^4,6,7^. HDACs can be divided into class I (HDAC1, 2, 3, and 8), class II (HDAC4, 5, 6, 7, 9, and 10), and class IV (HDAC11) based on sequence similarity. The class II HDACs are further divided into class IIa (HDAC4, 5, 7, and 9) and class IIb (HDAC6, and 10), according to their domain compositions. However, Class IIa HDACs lack any measurable enzymatic activity, and their deacetylase activity is minimal towards acetylated histones ^5^. Compared to Class I HDACs, Class IIa HDACs exhibit a prolonged N-terminal region that harbors a nuclear localization signal and a nuclear export signal, so they are found both in the nucleus and the cytoplasm ^5^. The presence of this unique N-terminal region suggests that Class IIa HDACs may participate in multi-protein complexes to exert their biological function. In human malignancies, Class I and II HDACs are considered oncoproteins that regulate gene expression to promote tumorigenesis and cancer development ^5–7^.

Specifically, HDAC7 has been implicated in the pathogenesis of colorectal^8,9^, pancreatic^10,11^, liver^12^, gastric^13^, lung^14^, breast^15^ and brain cancer^16,17^. Canonical histone proteins are incorporated on chromatin in a DNA replication dependent manner, while Histone H3.3 incorporates in a DNA replication independent manner ^18–20^. In Drosophila melanogaster and Arabidopsis thaliana, H3.3 is enriched in replication origins ^21–26^, while in human cells is associated with early replication and recruited on sites of DNA repair ^26–28^. Although H3.3 is mainly associated with euchromatin and regulation of active transcription ^20,29^, recently it was shown that it was also associated with heterochromatin on telomeres, pericentric heterochromatin and heterochromatin marks on gene regulatory regions ^29,30^. Association of H3.3 with euchromatin or heterochromatin is largely dependent on a group of proteins with chaperone activity that interact with H3.3 and deposit it on chromatin regions.

Euchromatin deposition is regulated by a HIRA/UBN1/CABIN1 chaperone complex while heterochromatin deposition of H3.3 depends on interaction with DAXX/ATRX ^31,32^. The balance between HIRA/UBN1/CABIN1 versus DAXX/ATRX deposition of H3.3 on euchromatin or heterochromatin respectively, may represent a crucial node in histone landscape reprogramming that enables complex cell fate decisions during tumorigenesis. Involvement of HDACs on these Histone chaperone complexes has only been suggested for the MIER1 complex but not for individual HDACs ^33^ and the participation of Class IIa HDACs in the regulation of the Histone H3.3 chromatin landscape has not been reported to date.

Here we show high expression of HDAC7 in glioblastoma tumor tissue from 84 patients, in patient-derived glioma stem cells (GSCs) from six patients and in pediatric diffuse pontine glioma (DIPG) cells from three patients. HDAC7 shows minimal histone deacetylase activity but forms protein-protein interactions with Histone H3.3 on chromatin. Specific inhibition of HDAC7 expression with an in vivo stable siRNA, results in an increase of heterochromatin mark H3K9me3 and heterochromatin spreading in cancer cells. Following HDAC7 inhibition, the interaction of H3.3 with HIRA is reduced, the association of H3.3 with DAXX and H3K9me3 is increased and H3.3 is deposited on H3K9me3^+^/DAPI^+^ heterochromatin nuclear foci. This results in increased spreading of H3K9me3 on the cancer genome and significant alteration of cancer cell gene expression. Inhibition of HDAC7 results in increase of DNA replication fork speed without affecting symmetry, resulting in replication stress and reduction of EdU incorporation. Finally, knockdown of HDAC7 expression induces significant sensitization of cancers cells to DNA damaging agents.

## Results

### HDAC7 is highly expressed in glioblastoma tumors and correlates with decreased patient survival

Using data from The Cancer Genome Atlas (TCGA), we compared the expression of all Class IIa HDACs in glioblastoma. HDAC-7 is the only Class IIa HDAC with significantly higher expression in glioblastoma as compared to the non-tumor samples (Figure 1A). Also, to show clinical correlation of HDAC7 expression in glioblastoma, we performed survival analysis on Class IIa high- and low-expressing patients. Patients were divided into these two groups based on the maximally ranked statistics using the log rank test as well as Gehan-Breslow-Wilcoxon method. High HDAC7 expressing patients show significantly lower probability of survival (p<0.004) (Figure 1B), while the expression level of other members of Class IIa HDACs has no significant correlation with patient survival.

**Figure 1:**
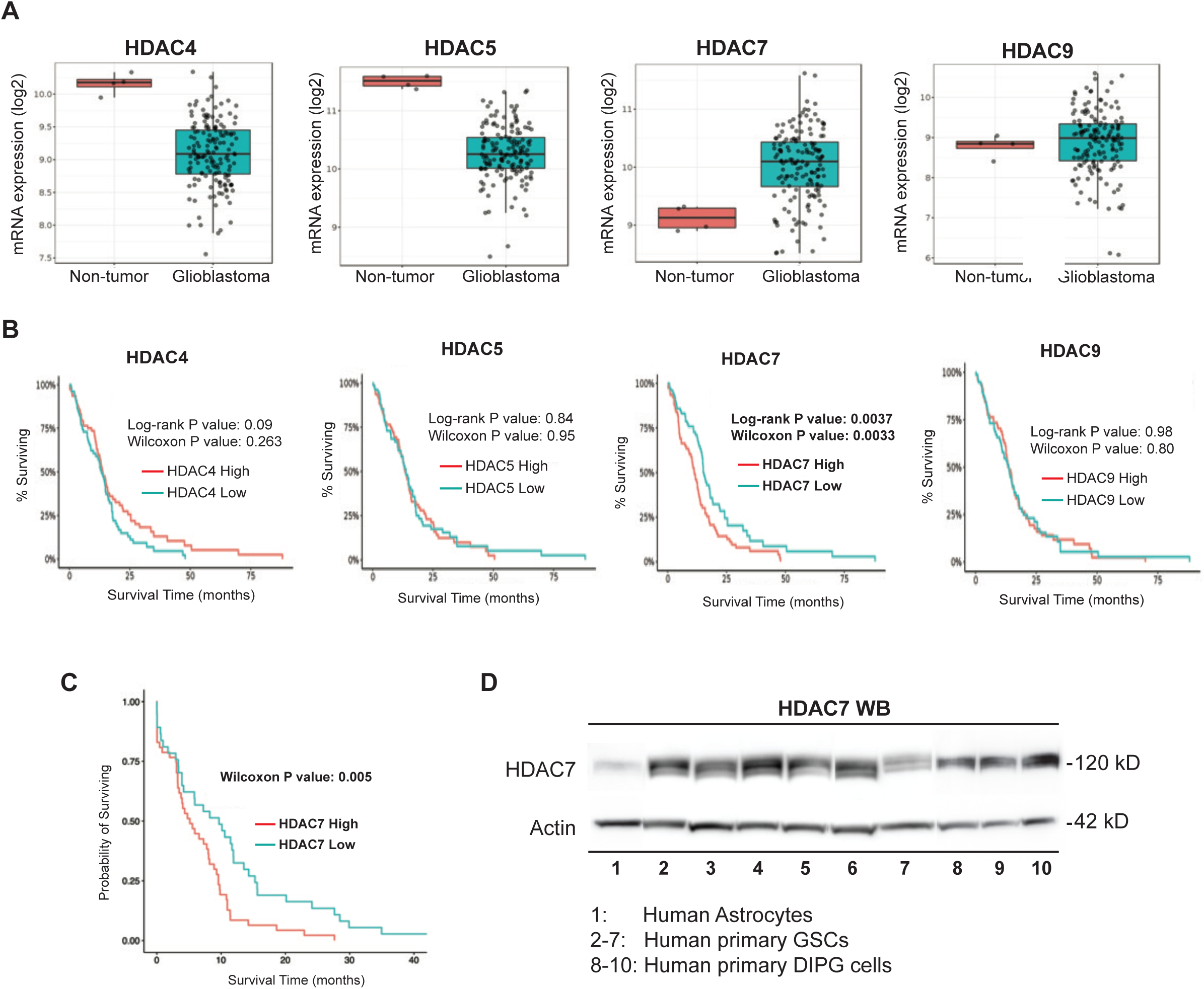
HDAC7 is highly expressed in glioblastoma tissue, GSCs and DIPG cells and correlates with significant decrease in patient survival. A) Transcript expression of Class IIa HDACs in glioblastoma compared to non-tumor control tissue using TCGA data. HDAC7 is the only Class IIa HDAC that is expressed significantly higher in glioblastoma compared to control tissue. B) Kaplan Meier survival analysis from the TCGA data, shows that high expression of HDAC7 is significantly (p<0.003) correlated with decreased patient survival. C) Kaplan Meier survival analysis in 84 patients with glioblastoma from Rhode Island Hospital reveals that high expression of HDAC7 correlates with significant decrease in survival probability (p<0.005). D) Expression of HDAC7 in 6 patient-derived GSCs and 3 patient-derived DIPG cells compared to normal human astrocytes as detected by Western blot (representative example of n=3 biological replicates). Actin was used as loading control.

In collaboration with the local Pathology Department at our hospital, we collected 84 formalin-fixed paraffin embedded (FFPE) tumor samples from patients with confirmed diagnosis of IDH WT Glioblastoma. We performed Nanostring gene expression analysis using the Pan Cancer Panel that contains 770 genes with confirmed roles in cancer. The expression data were clustered based on high versus low expression of HDACs. Classification of gene expression in HDAC7 “high” versus HDAC7 “low” expressing patients, showed that high HDAC7 expression correlates with significant increase in expression of 45 genes and inhibition of 2 genes. Functional classification of the significantly increased genes shows that several have known pro-tumorigenic roles like Vimentin, Lox, SERPINH1, TGFb, etc. (Supplementary Figure S1A). Finally, we performed Kaplan Meier survival analysis on the 84 patients based on high versus low expression of HDAC7 and showed that patients with high HDAC7 expression had significantly lower disease-free survival (Wilcoxon p<0.005) (Figure 1C). **Expression of HDAC7 in adult glioblastoma and pediatric DIPG cells.**

To determine expression levels of HDAC7 in human cancer cells, we performed Western blotting on lysates from 6 patients-derived GSCs with IDH WT glioblastoma and primary DIPG cells from three patients and compared expression to control human astrocytes. We show that HDAC7 is highly expressed in GSCs and DIPG cells while minimally expressed in control human astrocyte lysates (Figure 1D, representative example of three biological replicates).

The presence of nuclear localization and nuclear export signals makes HDAC7 capable of nuclear and cytoplasmic expression depending on the cell type ^34,35^. To determine the topology of HDAC7 expression in patient-derived GSCs, we performed immunofluorescent staining of HDAC7 expression (red) and Vimentin (green) as cytoplasmic marker. Nuclei were visualized with DAPI (blue) and z-stack images were acquired with a Zeiss Axiovert inverted microscope equipped with Apotome II. We show that HDAC7 is highly expressed in the nucleus of GSCs with minimal expression in the cytoplasm (Figure 2A).

**Figure 2:**
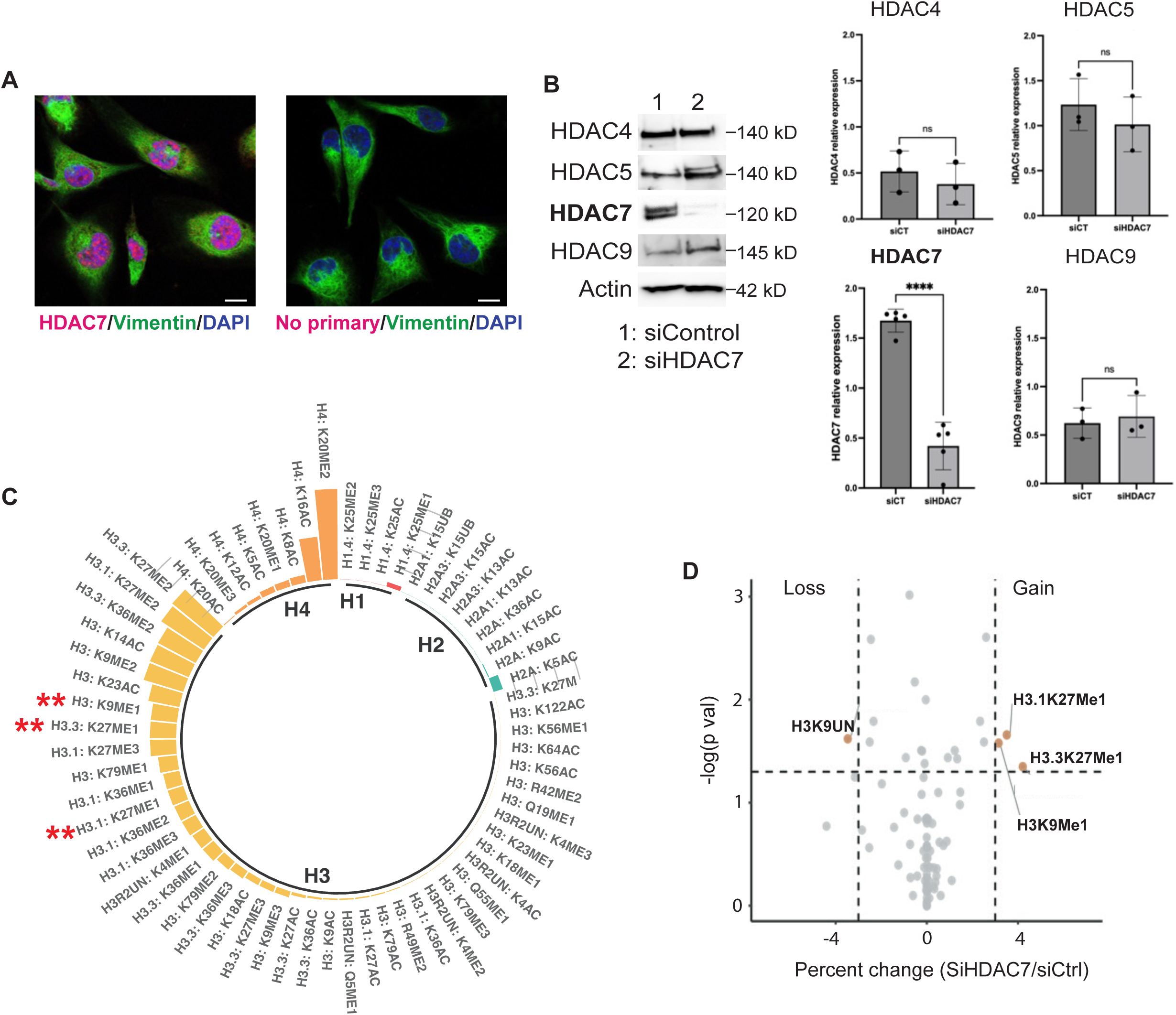
HDAC7 specific knock-down does not alter histone acetylation status but increases heterochromatin marks in GSCs. A) Nuclear expression of HDAC7 in patient derived GSCs stained with an HDAC7 antibody (magenta) and vimentin (green) as a cytoplasmic marker. No-primary antibody control shows lack of non-specific staining. B) Class specificity of the HDAC7 in vivo stable siRNA. siHDAC7 induces significant knockdown of HDAC7 protein expression (p<0.0001, n=5) while it does not affect the expression of the other members of the Class IIa HDAC family (n=3, n.s.). C) Circular plot shows percent abundance of histone modifications in GSCs. Asterisks are used to indicate histone sites undergoing significant alterations following HDAC7 knock down. D) Volcano plot of differentially modified histone marks in control versus siHDAC7 samples (cut-off of 3% change and P<0.05).

### Class specificity of HDAC7 siRNA

To discover a target specific siRNA for HDAC7, we designed, filtered through BLAST similarity search, and finally synthesized 4 individual siRNAs that target the open reading frame and 3’- untranslated region of human HDAC7. 50nM of each siRNA was separately transfected into patients derived GSCs and the expression of Class IIa HDACs was determined by Western blotting 72 hours after transfection. Control cells were transfected with non-targeting siRNA and Actin was used as loading control. The selected siRNA shows specific inhibition of HDAC7 protein expression (p<0.0001, n=5) and does not significantly affect the expression of HDAC4, HDAC5 and HDAC9 (Figure 2B and Supplementary Figure S1B & S1C).

### MS profiling of histone marks following inhibition of HDAC7 expression

To determine if HDAC7 has any effect on levels of histone modifications, we performed histone modification Mass Spectrometry (ModSpec-Active Motif) after inhibition of HDAC7 expression with our specific siRNA. Individual histone modifications were quantified by ModSpec and expressed as a proportion of all modified and unmodified residues (% of the peptide pool). At baseline (siControl GSCs), we found histones H3 and H4 to be highly modified. In contrast, H1 and H2 showed only minor to no modifications (Figure 2C). The most modified histone sites were H3K9, H3.1K27, H3.3K27 and H4K20 (Supplementary Figure S2). On aggregate, dimethylation of histone marks shows the highest abundance in comparison to other modifications in GSCs. To determine significant changes in levels of histone modifications following inhibition of HDAC7, we used a cut-off of 3% change and a P<0.05. Differences in the total enrichment of all marks (as a proportion of all modified and unmodified peptides) at each of the different histone sites are shown in Figure 2D. We show that inhibition of HDAC7 expression does not affect levels of histone acetylation suggesting minimal histone deacetylase activity but induces significant increase in H3K9me1 and reduction at levels of unmodified H3K9UN, which implies major chromatin landscape restructuring and increase of heterochromatin. In addition, we detect significant increase in H3.3K27me1 and H3.1K27me1, which is a mark that has been associated mainly with heterochromatin ^36,37^ but also with active transcription in embryonic stem cells ^38^.

### Rapid Immunoprecipitation Mass Spectrometry of Endogenous Proteins (RIME) to identify the HDAC7 interactome on chromatin

To interrogate how inhibition of HDAC7 results in increase of heterochromatin marks, we hypothesized that HDAC7 forms multiprotein interactions on euchromatin to participate in regulation of active transcription. We employed Rapid Immunoprecipitation Mass Spectrometry of Endogenous Proteins (RIME) to discover the HDAC7 protein-protein interaction network on chromatin of GSCs. GSCs (n=3) were fixed with formaldehyde to crosslink proteins and DNA that are in very close proximity. Nuclei were then isolated and HDAC7 immunoprecipitation was performed. The HDAC7 precipitated protein complex was subjected to Mass Spectrometry (RIME) which revealed 247 proteins constituting the HDAC7 interactome. The proteins were entered into STRING for network analysis using default settings and exported into Cytoscape for network construction. 229 from the 247 proteins formed a highly interconnected network (Figure 3A). To focus the analysis, we extracted the HDAC7 1^st^ degree interactome, which comprises the complete set of nodes and edges of proteins directly linked to HDAC7 (Figure 3B). The HDAC7 1^st^ degree interactome included 10 nodes, among them NCOR2, H3.3, H3.4, H3C13, H3C12, CTNNB1, CHD4, TBLX1XR1 and HSPA4, which together form a highly interconnected network (PPI p value = 5.28E-18). This network is enriched for transcription regulation pathways, notch signaling, chromatin modeling, and Beta-catenin signaling (Figure 3B). Next, we extended the analysis to include nodes connecting directly to the HDAC7 1^st^ degree interactome, thus generating the 1^st^ and 2^nd^ degree interactome (Figure 3C). The 1^st^ and 2^nd^ degree interactome was composed of 84 nodes that were significantly enriched for mitotic activity and cell cycle pathways (Figure 3C).

**Figure 3:**
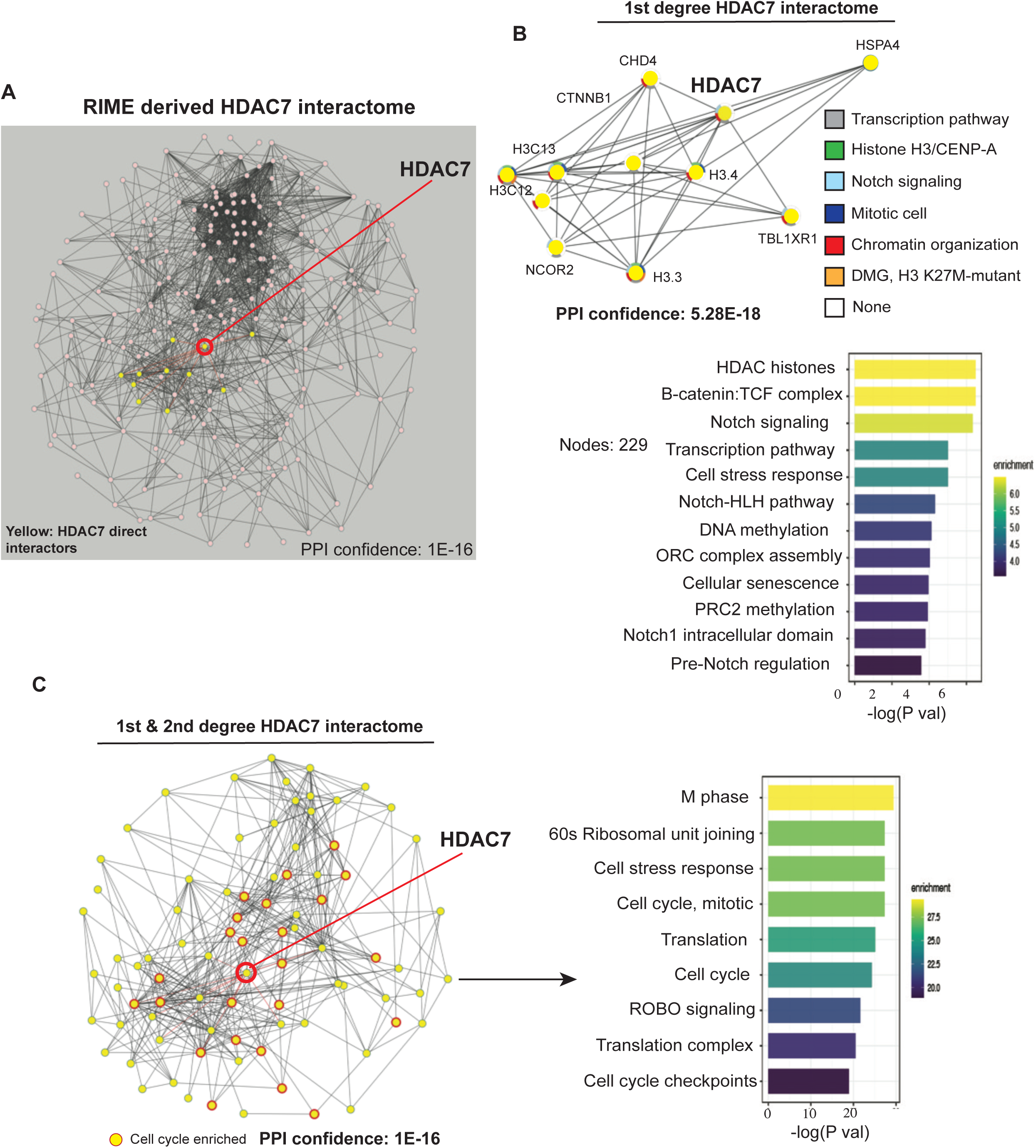
HDAC7 interactome on chromatin of patient derived GSCs. A) Network of 229 nodes derived from RIME identified proteins interacting with HDAC7 (1E-16). HDAC7 is shown near the center of the network. B) 1^st^ degree interactome: the proteins directly interacting with HDAC7. Each protein/gene is color-coded based on functional enrichment. Corresponding enrichment analysis shown. C) 1^st^ & 2^nd^ degree interactome: Proteins connected to those directly interacting to HDAC7. Corresponding enrichment analysis shown.

### HDAC7 interacts with Histone H3.3 and the H3.3 mutant variant H3K27M

To validate the interaction between HDAC7 and H3.3, we performed co-immunoprecipitation (co-IP) experiments using protein lysates from patients derived GSCs (n=3). Immunoprecipitation was performed with an HDAC7 validated antibody (Abcam) and the presence of H3.3 was detected with an H3.3. specific antibody (Active Motif). This showed that HDAC7 co-precipitates with H3.3 (Figure 4A) validating the RIME results. Since the H3.3 mutant variant H3K27M has been implicated in pediatric brain tumors, we performed co-IPs using lysates from patient derived DIPG cells (kind gift of Dr. Michelle Monje, Stanford University) and showed that HDAC7 interacts and co-precipitates with H3K27M (Figure 4B) suggesting a role for HDAC7 in these pediatric tumors.

**Figure 4:**
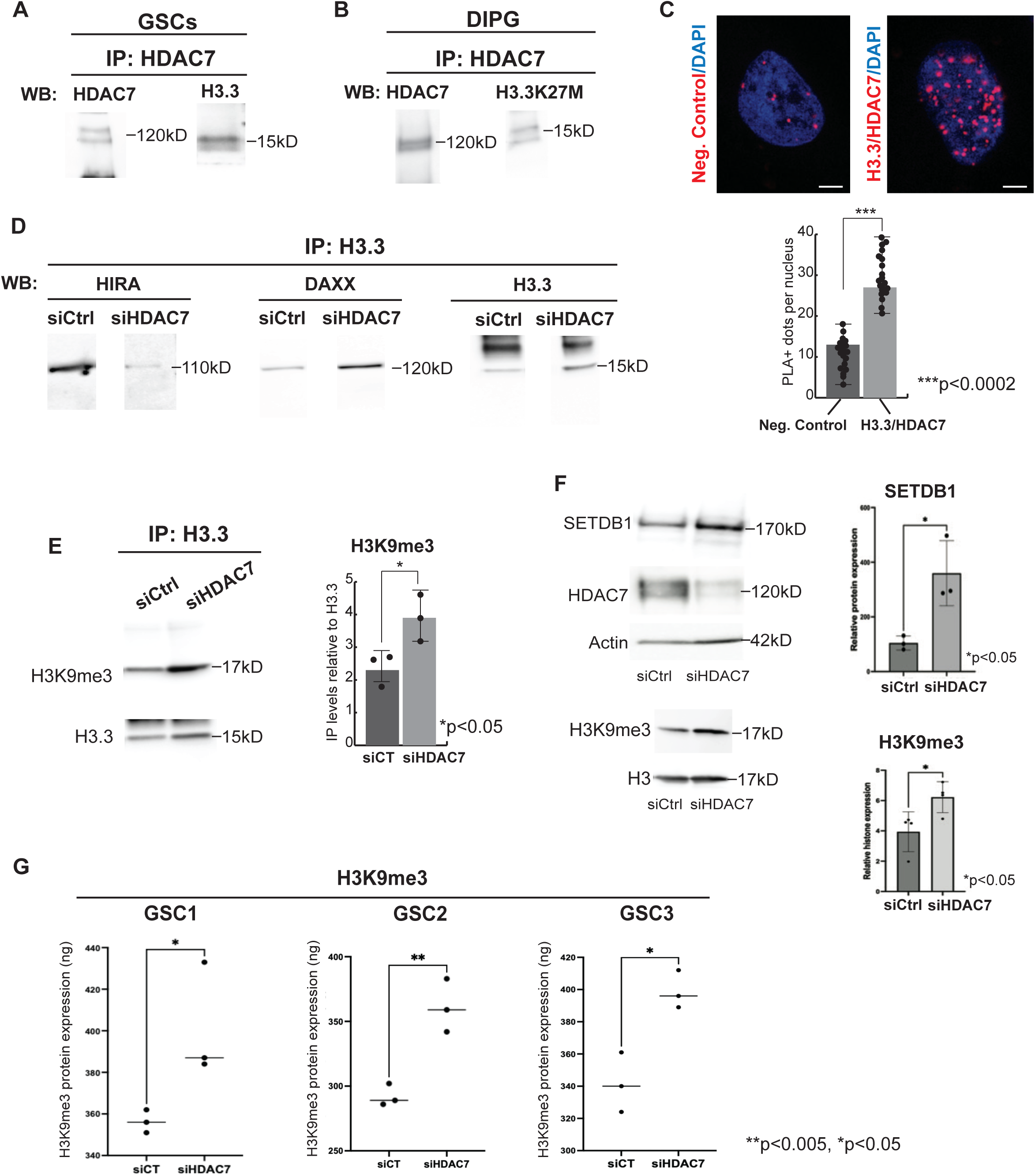
HDAC7 interacts with H3.3 and inhibition of HDAC7 reduces H3.3/HIRA interactions and increases H3.3/DAXX and H3.3/H3K9me3. A) HDAC7 and Histone H3.3 co-immunoprecipitation from nuclear lysates of GSCs. Representative example of 3 independent experiments. B) HDAC7 and oncohistone H3.3K27M co- immunoprecipitation from nuclear lysates of pediatric DIPG cells. Representative example of 3 independent experiments. C) Proximity Ligation Assay (PLA) for HDAC7 and H3.3 in GSCs shows specific interaction in the nucleus (red foci). Quantification was performed from 30 nuclei and the results are shown as number of PLA+ foci per nucleus (p<0.0002). D) Inhibition of HDAC7 results in inhibitions of H3.3 interaction with HIRA and increase in interaction of H3.3 with DAXX. Co-immunoprecipitations were repeated 3 independent times. E) HDAC7 knockdown results in significant increase in the association of histone H3.3 with the heterochromatin mark H3K9me3 (p<0.05, n=3). F) Inhibition of HDAC7 expression in GSCs results in significant increase in levels of SETDB1 and H3K9me3 (p<0.05, n=3, n=4). G) H3K9me3 ELISA on nuclear lysates from three independent patient-derived GSCs shows significant increase in H3K9me3 heterochromatin mark (*p<0.05, **p<0.005, n=3).

To verify the interaction between HDAC7 and Histone H3.3 in glioblastoma cells at a single molecule level, we performed proximity ligation assay (PLA) using the Duolink In Situ Red Starter Kit (Sigma). We show that H3.3 and HDAC7 specifically interact in the nucleus of glioblastoma cells (Figure 4C).

### Inhibition of HDAC7 reduces interaction of H3.3 with HIRA and increases interaction with DAXX and H3K9me3

To determine how HDAC7 affects the H3.3. landscape in cancer cells, we inhibited the expression of HDAC7 with siHDAC7 and analyzed the effects of HDAC7 inhibition on the interaction of H3.3 with its euchromatin chaperone HIRA, or its heterochromatin chaperone DAXX/ATRX. We show that inhibition of HDAC7 results in reduction of the interaction of H3.3 with HIRA in GSCs and increases the interaction of H3.3 with the heterochromatin chaperone DAXX (Figure 4D). These results suggest that inhibition of HDAC7 expression results in reprograming of the H3.3 landscape, reduction of H3.3. association with euchromatin chaperones and increased association of H3.3 with heterochromatin. To verify that inhibition of HDAC7 results in increased

association of H3.3 with heterochromatin, we performed co-IP of H3.3 with the heterochromatin mark H3K9me3. This shows that following inhibition of HDAC7 expression, the interaction of H3.3 with H3K9me3 is increased significantly (n=3, p<0.05, Figure 4E).

### HDAC7 inhibition results in increased levels of SETDB1, and H3K9me3

Since inhibition of HDAC7 results in increased association of H3.3. with heterochromatin, we asked whether siHDAC7 induces global increase of heterochromatin marks in cancer cells. We performed Western blot detection of H3K9me3 in patients derived GSCs (n=3) and show a significant increase of H3K9me3 (Figure 4F). In addition, protein levels of SETDB1, the histone methyltransferase that mediates the trimethylation of H3K9, was significantly elevated following inhibition of HDAC7 (Figure 4F). To verify the increase in levels of H3K9me3, we performed quantitative ELISA (Active Motif) to detect levels of H3K9me3 in patients derived GSCs (n=3) and again show significant increase following inhibition of HDAC7 (Figure 4G).

### Inhibition of HDAC7 induces heterochromatin spreading in cancer cells

To visualize and quantify the increased association of H3.3 with heterochromatin and the global increase of heterochromatin marks in cancer cell nuclei, we performed immunofluorescence localization of H3.3 and H3K9me3 with and without siRNA inhibition of HDAC7. Quantification of the number of H3.3 containing heterochromatin foci per nucleus from 30 individual cell nuclei shows that siHDAC7 induces a significant increase in localization of H3.3 within heterochromatin foci (Figure 5A-B). In addition, siHDAC7 induces significant increase in nuclear H3K9me3 expression (Figure 5C-D). To better visualize the spreading of heterochromatin within the cancer cell nucleus, we performed transmission electron microscopy (TEM) on patient derived GSC nuclei before and after inhibition of HDAC7 with siRNAs. We show that inhibition of HDAC7 results in spreading of electron-dense pericentric heterochromatin and increase of heterochromatin chromobodies (Figure 5E, arrows).

**Figure 5:**
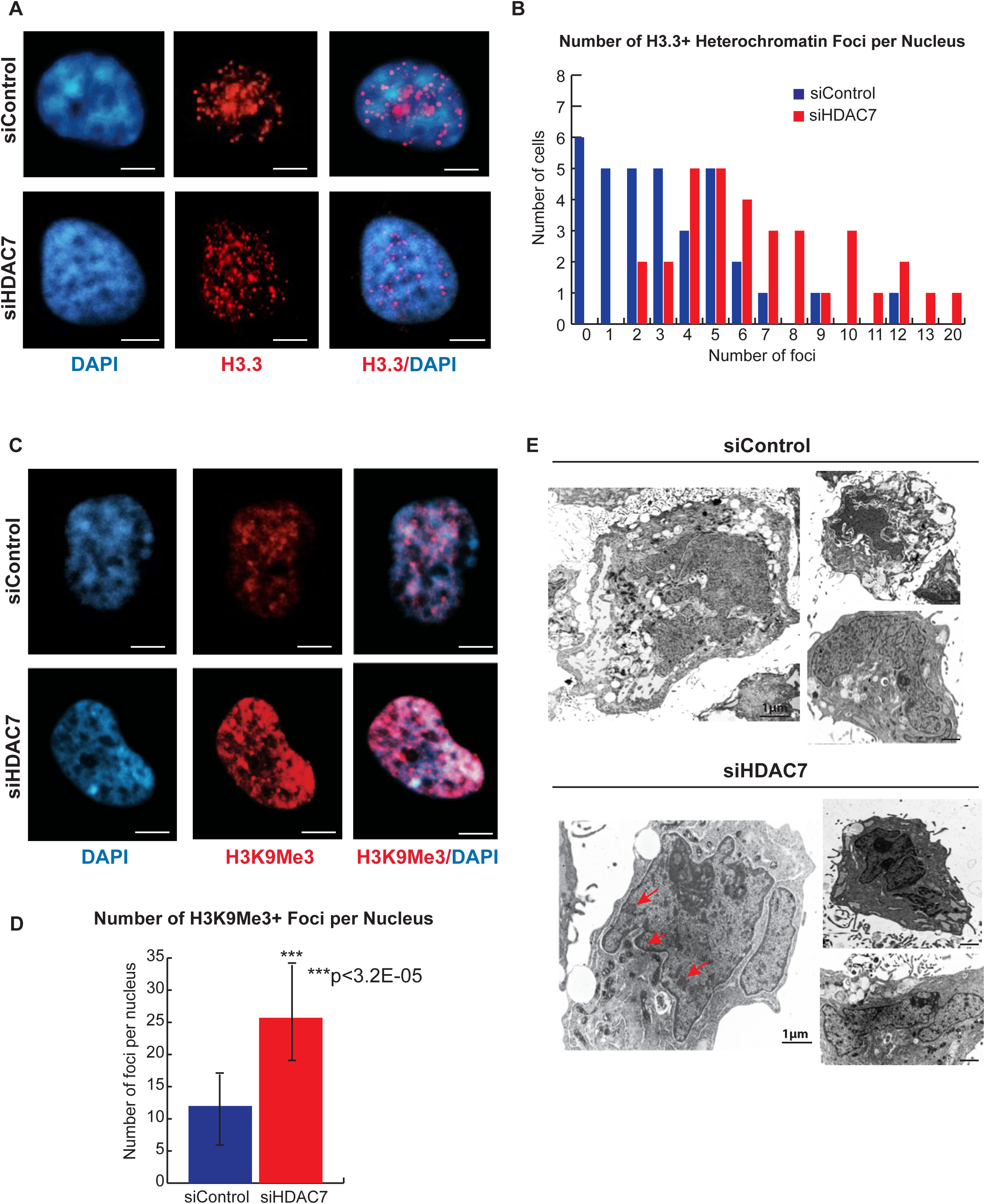
Inhibition of HDAC7 results in H3.3 landscape reprogramming and heterochromatin spreading in cancer cells. A) Immunofluorescence staining of H3.3 (red) and DAPI (blue) in GSCs before and after knock-down of HDAC7 shows that inhibition of HDAC7 expression results in redistribution of H3.3 from DAPI^(-)^ euchromatin nuclear domains to DAPI^(+)^ heterochromatin foci. B) Quantification of the number of H3.3+ heterochromatin foci per nucleus from 30 GSC nuclei per condition (siControl vs siHDAC7). There is a significant increase in the average number of H3.3+ heterochromatin foci following HDAC7 inhibition. Control average number of H3.3+ foci: 3.14 +/- 2.5, siHDAC7 average number of H3.3+ heterochromatin foci: 7 +/- 3.5. P<1.3E-05. C) Immunofluorescence staining of H3K9me3 (red) and DAPI (blue) in GSCs before and after knock-down of HDAC7 shows that inhibition of HDAC7 expression results in increase of the heterochromatin mark in the nucleus of GSCs. D) Quantification of the number of H3K9me3+ foci in the nucleus of 20 GSCs with and without inhibition of HDAC7 shows significant increase in the number of H3K9me3+ foci following inhibition of HDAC7 (p<3.2E-05). E) Transmission electron microscopy of patient derived GSCs before (siControl) and after (siHDAC7) inhibition of HDAC7 expression shows spreading of electron dense heterochromatin in pericentric nuclear domains (arrowhead) and nuclear chromobodies (arrows).

### Inhibition of HDAC7 results in increased H3K9me3 distribution in the cancer genome

To quantify the effect of HDAC7 inhibition in the spreading of H3K9me3+ heterochromatin on the cancer genome, we performed H3K9me3 ChIP-seq following transfection of GSCs with siControl or siHDAC7. To visualize the distribution profile of H3K9me3 we used ComputeMatrix and plotHeatmap (deepTools). Merged regions from siControl and siHDAC7 samples (n=3) were annotated then used for downstream differential read analysis to identify sites with increased H3K9me3. The signal distribution at the sites with increased H3K9me3 following inhibition of HDAC7, were obtained using the reference point mode. Regions were aligned based on the reference point, here the center was selected. Heatmap and distribution profile of the H3K9me3 was performed using plotHeatmap (deepTools) (Figure 6A). To identify sites with increased H3K9me3 signal, the log2FC of the reads between siControl and siHDAC7 KD samples was measured with log2FC of 1.5 as a cut-off. We identified 306 sites with significant increase in H3K9me3. Most of these sites annotated to gene-bodies (intronic sites) and intergenic sites. There were 214 genes in the vicinity of the 306 sites with increased H3K9me3 binding. Functional classification of these genes reveals genes with a role in proliferation, cell cycle regulation, DNA replication, glioma biology, ubiquitination and DNA damage (Figure 6B & Supplementary Table 1).

**Figure 6:**
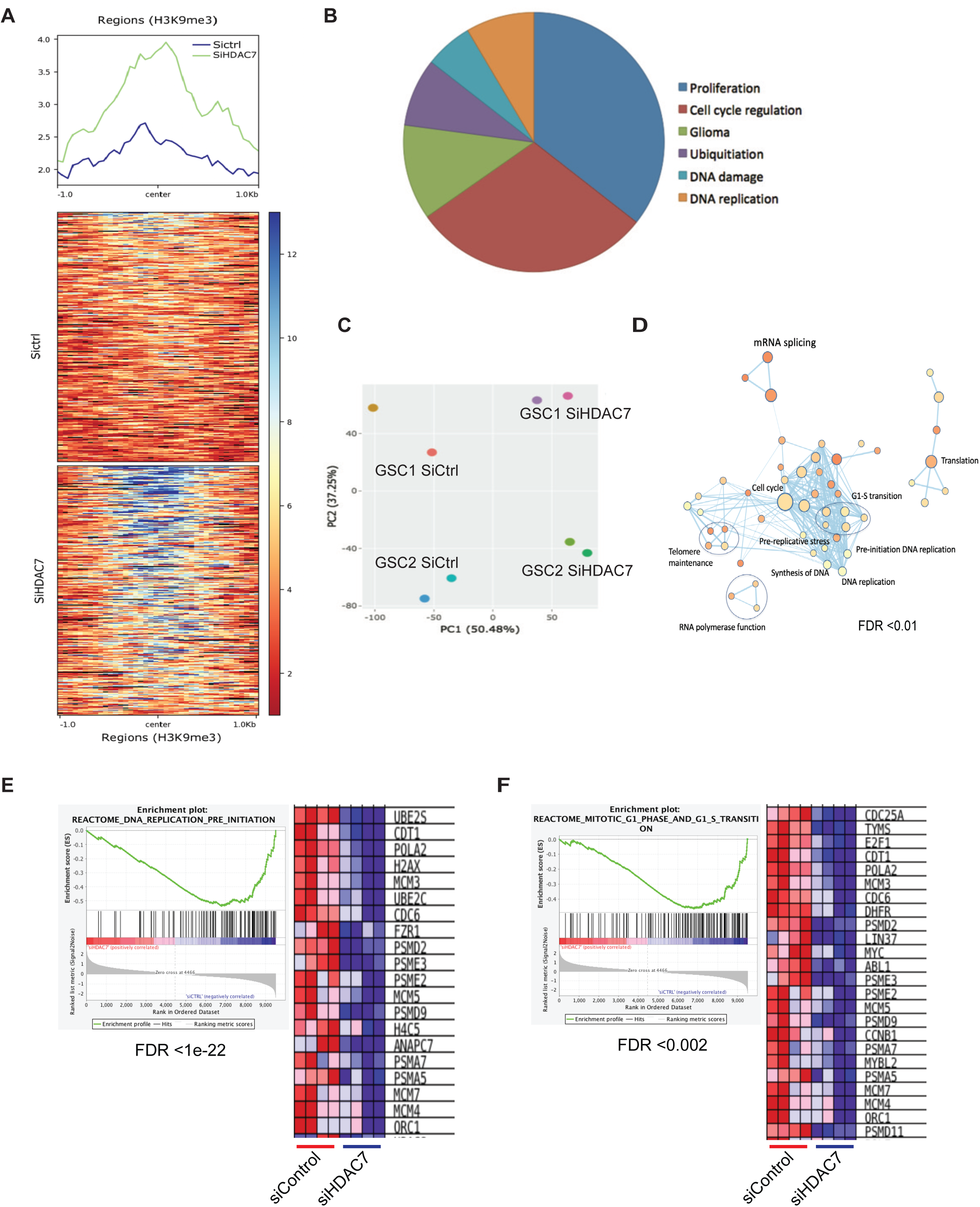
Transcript expression analysis and H3K9me3 ChIP-seq in GSCs following inhibition of HDAC7 expression. A) Read-count tag density aggregates of H3K9me3 profile (from three replicates). The signal distribution at the sites with increased H3K9me3 following inhibition of HDAC7, was obtained using the reference point mode. Regions were aligned based on the reference point, and the distribution of H3K9me3 is profiled using plotHeatmap relative to the center as a reference point. B) Pie chart depicting functional classification of genes in the vicinity of increased H3K9me3 signal following HDAC7 inhibition. Most prominent groups are shown as a percentage/fraction of queried/classified genes. C) PCA plot of GSCs in control versus siHDAC7 samples. D) Pathway enrichment for 194 downregulated genes (adj. p val < 0.001, FC>2) shows significant functional clustering for genes regulating cell cycle progression, G1-S transition, DNA replication and telomere maintenance. E & F) Gene Set Enrichment analysis shows that inhibition of HDAC7 induces significant decrease (FDR<1e-22, FDR<0.002) in major regulators of DNA replication (MCM3, MCM4, MCM5, MCM7, ORC1) and G1-S phase transition of cell cycle (MYC, E2F1, CDC6).

### Inhibition of HDAC7 results in broad transcriptomic changes in cancer cells

To assess the effect of HDAC7 inhibition at the transcript expression level, we performed RNA-seq with and without inhibition of HDAC7 expression in GSCs (n=2 biological replicates with n=2 technical replicates). PCA demonstrated appropriate segregation across conditions, suggesting that the siRNA targeting of HDAC7 induced reproducible changes across samples (Figure 6C). We identified 194 downregulated genes (adjusted p val < 0.001, FC>2) with significant functional clustering for genes regulating cell cycle progression, G1-S transition, DNA replication, telomere maintenance (Figure 6D). Gene Set Enrichment analysis (GSEA) shows that inhibition of HDAC7 induces significant decrease (FDR<1e-22, FDR<0.002) in major regulators of DNA replication (MCM3, MCM4, MCM5, MCM7, ORC1) and G1-S phase transition of cell cycle (MYC, E2F1, CDC6) (Figure 6E & 6F). In addition, genes with central role in cancer stem cell biology (Notch, Myc, Raf, JMJD1, etc.), embryonic stem cell signatures and FOXR2 target genes were downregulated following inhibition of HDAC7 expression (Supplementary Figure S3A & S3B).

To determine functional significance of the effect of HDAC7 inhibition in transcript expression and since transcripts of major regulators of cell cycle, DNA replication and cancer stemness were affected, we performed tumor-sphere formation assay with and without inhibition of HDAC7 in GSCs (n=3). Tumor-sphere formation provides a direct link to cancer stem cell self-renewal and division. We show that inhibition of HDAC7 results in significant inhibition of tumor sphere formation capabilities of human GSCs (p<0.0001) (Supplementary Figure S3C).

### Inhibition of HDAC7 alters cell cycle and DNA replication dynamics

To assess the effect of inhibition of HDAC7 on the rate of DNA replication in a per-cell manner, we quantified the incorporation of the thymidine analog 5-ethynyl-20-deoxyuridine (EdU) after a 20-min pulse of EdU in asynchronously growing GSCs (n=3). We determined EdU positive cells as a percent of total DAPI positive cells from 3 biological replicates, measuring 1000 nuclei from each experiment. We show that siHDAC7 results in significant inhibition of DNA synthesis as measured by EdU incorporation (Figures 7A-B, p<0.001). To examine if the same trend exists in DIPG cells we performed a 20-min pulse of EdU after siHDAC7 in patient-derived DIPG cells. In addition, we labelled the DIPG cells with H3K9me3 to determine if siHDAC7 results in increase of the heterochromatin mark. We show that siHDAC7 results in significant inhibition of EdU incorporation in DIPG cells (n=3, p<0.005) and significant increase in H3K9me3 positive nuclei (n=3, p<0.005) (Figure 7C). The same effects on DNA replication following HDAC7 depletion were also observed in the pancreatic cancer cell line PANC-1 (Supplementary Figure S3D).

**Figure 7:**
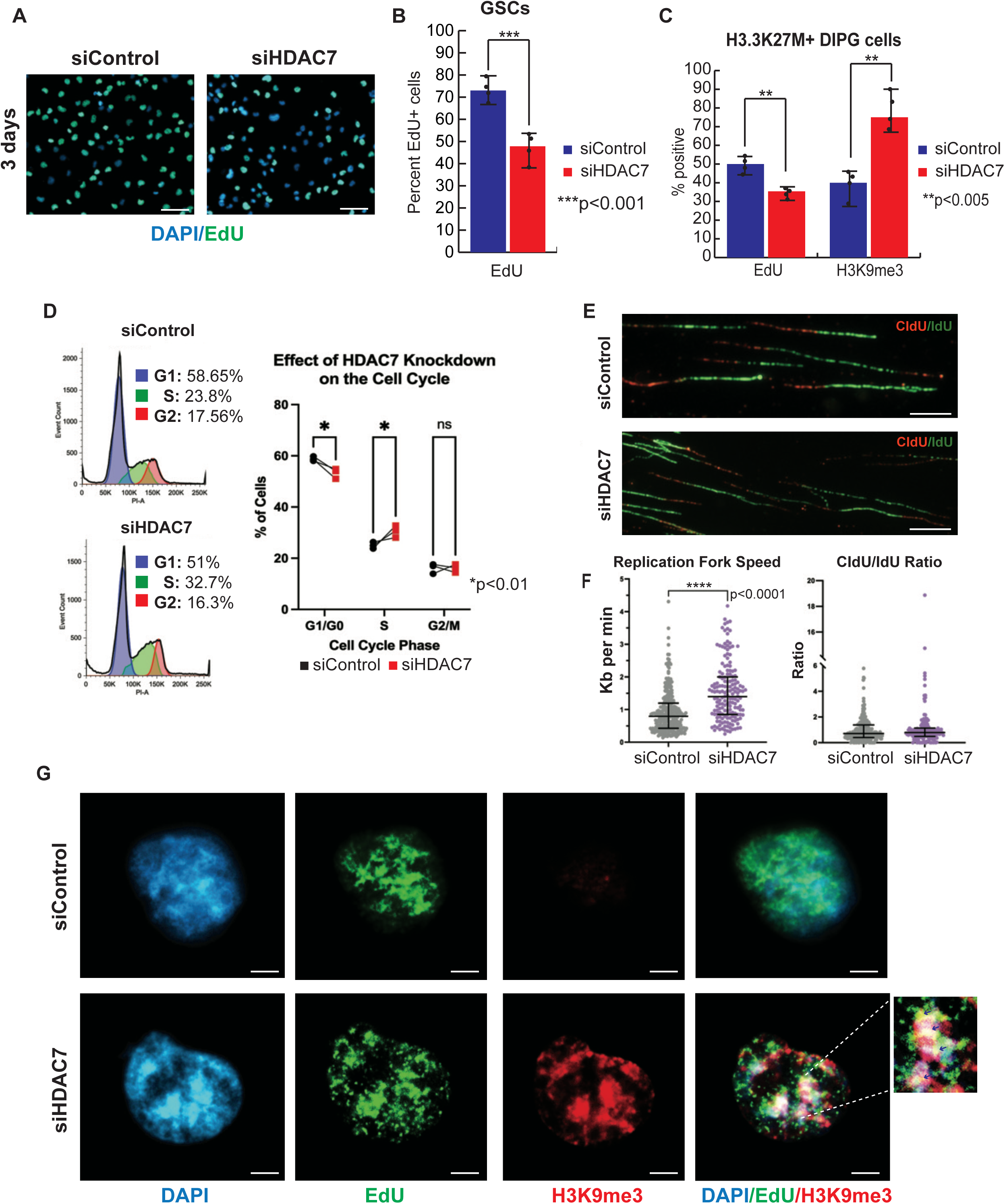
Inhibition of HDAC7 affects DNA synthesis and replication dynamics in cancer cells. A) Immunofluorescence staining of EdU incorporation 3 days after inhibition of HDAC7 expression with siHDAC7 in human GSCs. EdU is stained green while DAPI positive nuclei are blue. Magnification: 20X. B) Quantification of EdU positive cells as a percent of DAPI positive nuclei shows that siHADC7 results in significant (p<0.001) inhibition of EdU incorporation. Quantification was performed from 3 independent experiments counting 1000 DAPI positive nuclei. C) siHDAC7 results in significant inhibition of EdU incorporation and significant increase in H3K9me3 positive nuclei in human primary H3.3K27M+ DIPG cells (p<0.005). Results were quantified from 3 independent experiments. D) Inhibition of HDAC7 results in significant reduction of cells in G1 phase and significant increase of cells in S phase (p<0.01). Experiments were repeated in three biological replicates. E) Representative images of CIdu/IdU staining of DNA fibers from siControl and siHDAC7 GSCs. Magnification: 60X. F) Quantification of replication speed from n=291 siControl and n=167 siHDAC7 fibers shows that inhibition of HDAC7 results in significant increase in replication fork speed (p<0.0001, Mann-Whitney U test), while fork symmetry as defined by the ratio of CIdU/IdU is not affected. G) Immunofluorescence staining of EdU and H3K9me3 in siControl and siHDAC7 cells. EdU is stained green, H3K9me3 red and DAPI blue. Inhibition of HDAC7 results in increase of H3K9me3 in the nucleus and co-localization of H3K9me3 within EdU positive nuclear foci (right panel and higher magnification inlet).

Since depletion of HDAC7 reduces DNA synthesis, we examined the effects of siHDAC7 on cell cycle progression. We show that HDAC7 knockdown results in significant reduction of cells transitioning through the G1 phase and a significant increase in the number of S phase cells without any change in G2 phase cells (Figure 7D). Given the inhibition of DNA synthesis following depletion of HDAC7 (Figures 7A-C), the increase in S-phase cells was surprising and prompted further investigation into the role of HDAC7 depletion in replication dynamics. We performed single molecule DNA fiber analysis (Figure 7E) to determine effects of HDAC7 depletion in replication fork speed and symmetry. We show that HDAC7 knockdown results in significant increase of replication fork speed without affecting fork symmetry (Figure 7F) suggesting that the replication machinery may react to the increase in heterochromatin spreading following HDAC7 depletion by increasing fork speed to overcome heterochromatinization. To determine if H3K9me3 heterochromatin mark localizes on EdU positive replication foci we performed immunofluorescence staining of GSC nuclei with H3K9me3 following EdU incorporation in siHDAC7 and siControl cells. We show that inhibition of HDAC7 expression results in co-localization of the heterochromatin mark H3K9me3 with EdU positive replicating foci (Figure 7G).

Inhibition of HDAC7 has been associated with senescence in aging dermal fibroblasts and in certain cancer cell lines^39,40^. In our data, inhibition of HDAC7 expression does not induce a senescence gene expression signature (Supplementary Figure S4A). In addition, the number of cells entering the G0 and G2 phase of the cell cycle is not different between siControl and siHDAC7 cells and the overall cell cycle profile is not showing senescent cell cycling characteristics like G1 arrest and low S-phase entry (Figure 7D). Finally, β-galactosidase staining of siControl and siHDAC7 cells shows no significant difference (Supplementary Figure S4B & S4C). Based on these data, we conclude that depletion of HDAC7 does not result in a senescent phenotype in GSCs.

### HDAC7 inhibition results in replication stress, DSBs and sensitization of cancer cells to DNA damaging agents

It has been reported that cells can effectively buffer changes in fork speed but if velocity increases above a 40%, genomic integrity is compromised^41^. In our data, the median fork speed in siControl cells is 0.79 kb/min, while the median for the siHDAC7 cells is 1.4 kb/min which corresponds to a 57% gain of replication speed following depletion of HDAC7 (Figure 7F). In addition, siHDAC7 results in significant inhibition of MCM 2-7 expression (Figure 8A) suggesting reduction in the number of active replication origins. To compensate, the remaining forks may move faster to complete genome duplication within S-phase. We examined if the significantly increased fork speed results in replication stress and DSBs. We quantified phosphorylation of RPA2 and levels of γH2AX and BRCA2 in siControl and siHDAC7 cells. We show increased phosphorylation of RPA32-S4/8 without change in levels of γH2AX, while BRCA2 expression was lower after HDAC7 depletion (Figure 8B). These data suggest that inhibition of HDAC7 expression in GSCs results in replication stress, while reduction of BRCA2 expression suggests that homologous recombination (HR) repair to protect replication forks is impaired following inhibition of HDAC7 expression. This creates a vulnerable replication environment that could further sensitize the cancer cells to DNA damaging agents. To demonstrate this, we incubated siControl and siHDAC7 GSCs with 150uM of the alkylating agent Temozolomide (TMZ) and show that addition of TMZ in siHDAC7 treated cells results in increase of γH2AX expression (Figure 8C). To show if this results in growth inhibition of cancer cells we incubated siControl and siHDAC7 treated cells with 150uM or 300uM of TMZ and show that addition of TMZ to siHDAC7 treated GSCs, results in significant growth inhibition (n=3 biological replicates, *p<0.01, ***p<0.001) compared to GSCs treated with TMZ alone and the effect is concentration dependent (Figure 8D). Similar effects on induction of chromatin bound γH2AX expression and growth inhibition of cancer cells were noted following treatment of GSCs (n=3) with 50uM of Etoposide (Figure 8E) and Camptothecin respectively (Supplementary Figure 4D).

**Figure 8:**
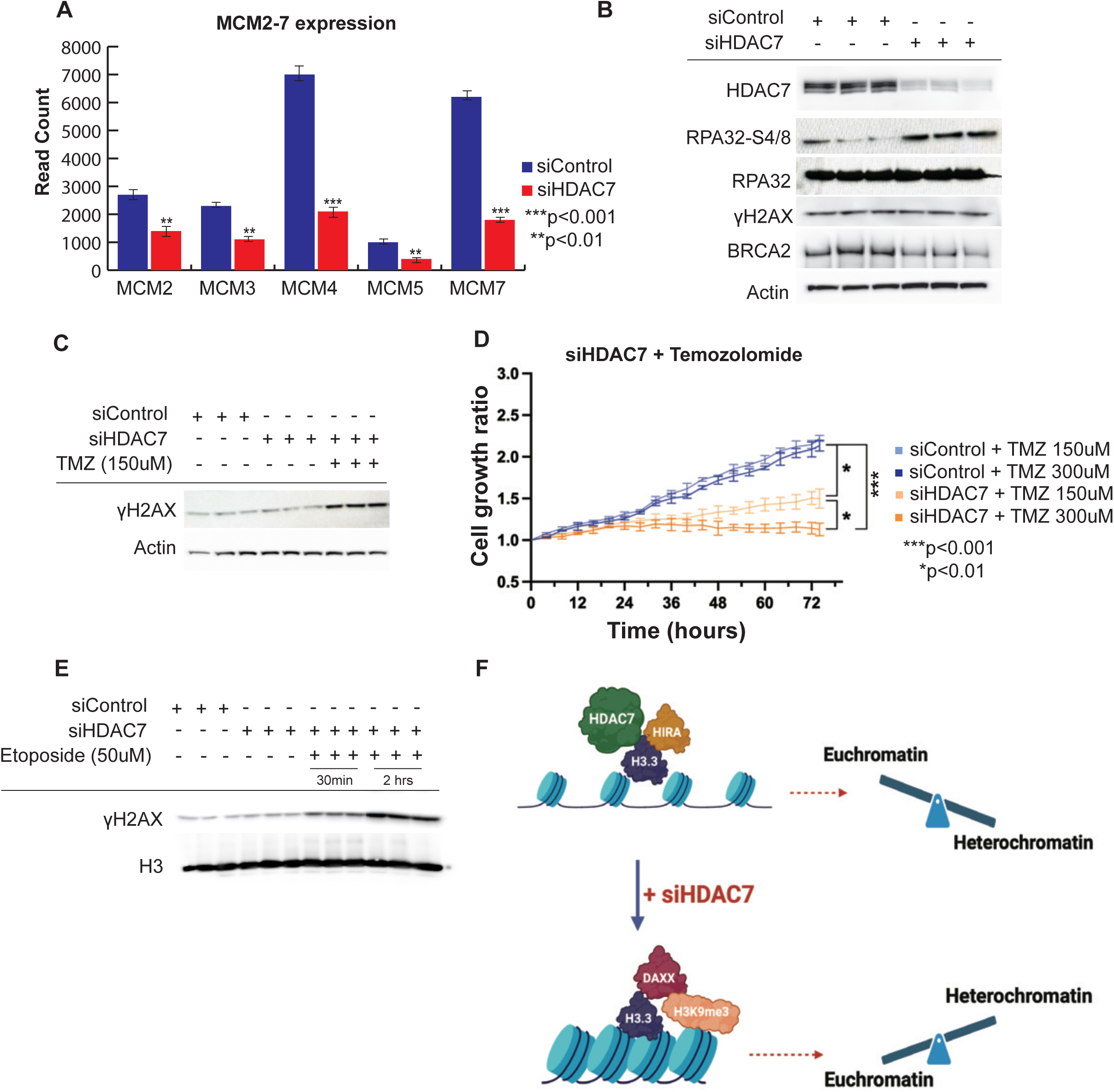
Inhibition of HDAC7 induces DNA replication stress and sensitizes cancer cells to DNA damaging agents. A) RNA-seq expression analysis of MCM 2-7 following inhibition of HDAC7 shows significant reduction of MCM 2, 3, 4, 5, 7 expression (***p,0.001, **p<0.01). B) Inhibition of HDAC7 results in increase of RPA32- S4/8 expression, reduction of BRCA2 expression while γH2AX is not affected. Western blots were performed in three independent samples and total RPA32 and Actin are used as loading controls. C) Addition of 150uM of TMZ in siHDAC7 treated samples results in increase of γH2AX expression (n=3). D) Real-time quantification of cell growth using Incucyte, in cells treated with TMZ alone (150uM or 300uM) or TMZ and siHDAC7. Inhibition of HDAC7 in combination with TMZ results in significant inhibition of cellular growth compared to TMZ alone in a concentration dependent manner (n=3 per condition, *p<0.01, ***p<0.001). E) Addition of 50uM Etoposide in siHDAC7 treated cells results in increase of chromatin-bound γH2AX expression (n=3). Histone H3 was used as loading control. F) Model depicting the role of HDAC7 in chromatin dynamics. HDAC7 binds to H3.3 and HIRA complexes favoring a euchromatic chromatin environment. Inhibition of HDAC7, results in deposition of H3.3 on heterochromatin in complex with DAXX and H3K9me3 resulting in dynamic shifting of the chromatin balance towards heterochromatin and epigenetic restriction.

## Discussion

Class IIa HDACs have limited catalytic activity in eukaryotic cells primarily due to a substitution of a tyrosine with histidine within the enzymatic pocket of these molecules ^42^. Although Class IIa HDACs can bind to acetylated lysine, they don’t exert deacetylase activity and it has been proposed that they function to attract other HDAC subtypes to deacetylate the substrates^43,44^. Several studies have highlighted the role of Class IIa HDACs in cancer, with the exact function of each member varying depending on the type of cancer^45^. HDAC7 has been implicated in the regulation of cancer stem cell fate, specifically in breast and ovarian tumors where it was shown that overexpression of HDAC7 can increase the cancer stem cell phenotype ^46^. HDAC7 binds near transcription start sites (TSS) and super-enhancers (SEs) of oncogenes in breast cancer stem cells and contributes to their transcriptional regulation. However, the mechanism by which HDAC7 maintains levels of H3K27Ac at TSS and SEs remains unclear ^45,47^.

Our findings provide compelling evidence supporting the interaction between HDAC7 and Histone H3.3 in glioblastoma cells. Through RIME, co-immunoprecipitation and PLA, we demonstrated a specific interaction between these proteins. This adds to the growing body of literature elucidating the complex interplay between histone molecules and their regulatory enzymes. The interaction between HDAC7 and Histone H3.3 suggests a role for HDAC7 in chromatin remodeling and gene regulation.

The specific interaction with H3.3, a histone variant associated mainly with active chromatin states, underscores the importance of HDAC7 in maintaining chromatin homeostasis. Histone H3.3 is deposited into euchromatin in a DNA replication independent manner. Specifically, H3.3 forms complexes with the histone chaperone HIRA to mediate DNA synthesis independent nucleosome assembly ^48^. Our study reveals that HDAC7 actively participates in these euchromatin complexes by interacting with H3.3 and HIRA. Inhibition of HDAC7 leads to a reduction in the association of H3.3 with HIRA and an increase in its association with DAXX and H3K9me3. Consequently, the H3.3 landscape undergoes remodeling in cancer cells, resulting in the spread of heterochromatin and its association with heterochromatin foci. Our findings unveil a novel aspect of HDAC7 biology in cancer. Notably, the observed increase in heterochromatin upon HDAC7 inhibition suggests a potential role for HDAC7 in maintaining the delicate balance between euchromatin and heterochromatin, which is often disrupted in cancer cells (Model, Figure 8F).

The heterochromatin histone mark H3K9me3 is associated with transcriptionally repressive regions of the genome and is a hallmark of constitutive heterochromatin, often found at pericentromeric and telomeric regions ^49^. As cells replicate, histone marks like H3K9me3 are diluted due to the incorporation of new, unmodified histones.

The co-localization of EdU (a marker for DNA synthesis) with H3K9me3 after inhibition of HDAC7 expression suggests that even in the context of active DNA replication, mechanisms are in place to ensure that heterochromatin marks are transmitted to daughter cells, supporting the inheritance of epigenetic states and stable gene repression.

Our data indicate that siHDAC7 treatment leads to a notable disruption of cell cycle progression, specifically by inhibiting the transition through the G1 phase and inducing a significant increase in the proportion of cells in the S phase, suggesting an accumulation of cells attempting to initiate or continue DNA replication. However, despite more cells entering the S phase, they do not seem to progress through it efficiently, as evidenced by the lack of accumulation in G2 phase cells and the significant reduction on cells actively incorporating EdU. This could imply that cells are spending more time in the S phase, potentially due to replication stress or inefficiencies in DNA replication. DNA fiber analysis reveals significant increase of replication fork speed, which may diminish the fidelity of DNA polymerases, potentially contributing to genomic instability^41,50^. At the same time, the reduction in MCM 2-7 expression could explain the increased fork speed^51^ as the cells are compensating the reduced number of active forks by increasing the speed of replication. Reduced MCM levels and insufficient origin firing might result in fewer global replication stress signals. Notably, the reduction in BRCA2 expression could also contribute to the increase in replication speed as shown in PARPi treated BRCA1-deficient cells^41^. Eventually, the increase in fork speed in conjunction with inhibition of MCM 2-7 and reduction in BRAC2 expression after depletion of HDAC7, result in genomic instability and replication stress. This DNA replication environment is particularly vulnerable to additional stress by DNA damaging agents. In fact, TMZ and Etoposide induced lesions appear to overwhelm the repair capacity of the cells, leading to significant DNA damage and increase in γH2AX expression. This highlights a potential therapeutic opportunity for combining inhibition of HDAC7 as a replication stress-inducing strategy with DNA damaging agents.

The identification of HDAC7 as a regulator of histone H3.3 chromatin dynamics and DNA replication offers a rationale for exploring the potential therapeutic benefits of inhibiting HDAC7 in cancer. By promoting heterochromatin spreading, HDAC7 inhibition may suppress oncogenic transcriptional programs, induce replication stress, and sensitize cancer cells to DNA-damaging agents, ultimately leading to cancer cell death. Further research in this field promises to reveal new avenues for precision medicine by therapeutic induction of epigenetic restriction in cancer.

## STAR Methods

### Primary glioblastoma stem cell isolation and culture

The institutional review board (IRB) of Rhode Island Hospital has approved the collection of patient-derived glioblastoma tissue. All collections were performed with written informed consent from patients in completely de-identified manner and the studies were performed in accordance to recognized ethical guidelines (Belmont Report). Primary GSC spheres were cultured from human glioma samples as previously described^52^. GSCs used in this study were authenticated by ATCC using short tandem repeat analysis. GSCs used were between passages 5 and 30 and cultured either as spheres or as attached on fibronectin-coated plates (10 mg/mL, Millipore Sigma) in a medium of 1X Neurobasal Medium, B27 serum-free supplement, minus Vitamin A, 100X Glutamax (Fisher Scientific), 1 mg/mL Heparin (STEMCELL Technologies), 20 ng/mL epidermal growth factor (Peprotech), 20 ng/mL bFGF (Peprotech). All GSC cultures are routinely tested for Mycoplasma contamination using the MycoSensor qPCR assay (Agilent).

### Analysis of publicly available cancer datasets

For survival analyses, we used the Cancer Genome Atlas (TCGA) patient cohort. Patients were assigned to high/low-risk groups based on HDAC7 expression levels. The datasets were downloaded from GlioVis and visualized in the Kaplan-Meier Plotter^53^, GlioVis^54^ web tools, and via the R Bioconductor package.

### Nanostring PanCancer progression analysis

We collected RNA from 84 formalin-fixed paraffin-embedded samples from glioblastoma patients and performed gene expression analysis using the Nanostring PanCancer progression panel that comprises the vital components of pathways involved in the complex interplay between the tumor and its microenvironment. Overall survival was estimated by the Kaplan-Meier method at GraphPad prism. Pathway enrichment and Gene ontology enrichment analyses were performed on the differentially expressed genes using R package DESeq^55^.

### siRNA transfection

GSCs were grown over fibronectin/HBSS-treated culture plates in antibiotic-free media supplemented with Neurobasal-A, B-27-A, Glutamax, growth factors (EGF and FGF), and heparin, and GSCs were incubated at 37°C with 5 % CO2 overnight. 50nM of HDAC7 siRNAs was used for all experiments. The sequences of the 4 HDAC7 siRNAs used in this study and the control non targeting siRNA are:

siRNA-1 sense: 5’ G.A.C.A.A.G.A.G.C.A.A.G.C.G.A.A.G.U.G 3’

antisense: 5’ C.A.C.U.U.C.G.C.U.U.G.C.U.C.U.U.G.U.C 3’

siRNA-2 sense: 5’ G.C.A.G.A.U.A.C.C.C.U.C.G.G.C.U.G.A.A 3’

antisense: 5’ U.U.C.A.G.C.C.G.A.G.G.G.U.A.U.C.U.G.C 3’

siRNA-3 sense: 5’ G.G.U.G.A.G.G.G.C.U.U.C.A.A.U.G.U.C.A 3’

antisense: 5’ U.G.A.C.A.U.U.G.A.A.G.C.C.C.U.C.A.C.C 3’

siRNA-4 sense: 5’ U.G.G.C.U.G.C.U.U.C.U.C.G.G.G.U.U.A.A 3’

antisense: 5’ U.U.A.A.C.C.C.G.A.G.A.A.G.C.A.G.C.C.A 3’

Non-targeting control siRNA- sense: 5’ A.G.C.G.A.C.U.A.A.A.C.A.C.A.U.C.A.A 3’

antisense: 5’ U.U.G.A.U.G.U.G.U.U.U.A.G.U.C.G.C.U 3’

### Tumor sphere formation assay

Tumor sphere formation assay was performed as described before^56^. Sphere formation by visual counting was estimated after seven days in a blinded, unbiased format by counting the number of spheres in each well by two independent researchers.

### RNA Sequencing

Next-generation RNA-sequencing was performed using an Illumina HiSeq2500 system (Azenta). Sequence reads were aligned to the human genome using *gsnap*. Genomic locations of genes and exons were extracted from the *refGene*.*txt* file (http://hgdownload.cse.ucsc. edu/goldenPath/hg38/database/refGene.txt.gz). Fo analysis, we used a false discovery rate FDR<0.001.

### Mass Spectrometry for Histone Modifications (Mod-Spec)

Five million GSCs were harvested, washed twice with PBS and snap frozen. Successful knock-down of HDAC7 was confirmed by Western blot and the samples were sent to Active Motif to perform Mod Spec per their in-house protocols. The entire Mass Spec process was repeated three separate times for each sample. The data was quantified using Skyline^57^ and presented as the percent of each modification within the total pool of that tryptic peptide.

### Rapid-immunoprecipitation-mass-spectrometry-of-endogenous-protein (RIME)

Two biological replicates of GSCs (5 x 10^8^ cells per replicate) were fixed for 8 minutes with a formaldehyde solution of 11% Methanol Free Formaldehyde, 0.1M NaCl (Invitrogen™), 1 mM EDTA, pH 8 (Invitrogen™), and 50mM HEPES, pH 7.9 (Fisher BioReagents™). Fixation was stopped using 2.5 M Glycine solution in water (Sigma). Fixed cells were washed with chilled PBS-Igepal (0.5% Igepal CA-630 in PBS) and PMSF (Active Motif). Washed cells were centrifuged, pelleted, and snap-frozen on dry ice and sent to Active Motif for RIME® service^58^. PEAKS Studio was used for visualization and analysis, and the “Decoy Fused Method” was applied for calculating FDR where the default cutoff is set to -10logP > 20 to ensure high-quality peptide spectrum matches and increase confidence in the proteins list obtained.

### Co-immunoprecipitation followed by western blot

After the respective course of treatment, GSCs (10 million cells / Co-IP) were collected and prepared for nuclear extraction and purification using a nuclear extraction kit (Active Motif). Nuclear extracts were treated with an enzymatic shearing Cocktail for DNA digestion to release undissociated protein complexes, including histone proteins and DNA-binding proteins from the DNA. 500 μg of extracted and purified nuclear lysate was used per IP per after pre-clearing with normal IgG (same concentration as the respective IP antibody) for 1 hour on a rotator at 4°C. Pre-cleared nuclear lysates were then combined with 25 μl of Protein G beads and incubated for 1 hour on a rotator at 4°C to clear away any IgG nonspecific binding. After separating the magnetic beads on a magnetic stand, the collected nuclear supernatant was then combined with the IP antibodies of either 1 μg HDAC7 (Cell Signaling) or 4 μg H3.3 (Abcam) and incubated for 4 hours on a rotator at 4°C. The immunoprecipitated samples were then combined with 25 μl of Protein G beads and incubated for 1 hour on a rotator at 4°C. After separating the magnetic beads on a magnetic stand, the immunoprecipitated samples (now bound to the beads) were washed four times with Co-IP wash buffer (Active Motif). The washed immunoprecipitated samples bound to beads were then resuspended in 20 μl of 2X Reducing Loading Buffer (130 mM Tris pH 6.8, 4% SDS, 0.02% Bromophenol blue, 20% glycerol, 100 mM DTT) and boiled at 95°C for 5 minutes. Magnetic beads were then separated from desaturated samples using a magnetic stand. All extraction and IP steps were performed in buffers supplemented with Protease and Phosphatase and Inhibitors to limit further protein modifications (expression, proteolysis, dephosphorylation, etc.). Immunoprecipitated samples were analyzed by Western blot. **Histone lysates extraction and purification**

The histone acid precipitation technique was used to extract pure crude histones from GSCs following the transfection treatment. After the course of treatment, GSCs were detached using StemPro Accutase (Gibco), and spun-down pellets were collected.

Pellets were then resuspended in the appropriate volume of ice-cold histone extraction buffer (Active Motif) and incubated overnight at 4°C on a rotator. On the next day, cell suspensions were centrifuged to collect core histone extracts. Acid extractions were pH measured and neutralized with the needed volume of 5X neutralization buffer until it reached complete neutralization. Core histones were then purified with column elution, and purified histone lysates were stored at -80°C for downstream experiments.

### H3K9Me3 ELISA

Purified core histone extracts prepared as described above were quantified for histone lysine trimethylation at lysine 9 following ELISA protocols from Active motif (Catalogue # 53109). Absorbance at 450nm with a reference wavelength of 655 nm was read on Promega™ GloMax® Plate Reader. Absorbance was normalized after subtracting blank wells. The quantification for methyl Lys9 from histone H3 in the samples was obtained using a standard curve plotted in GraphPad prism.

### Electron microscopy (EM)

Treated GSCs were harvested, and pellets were resuspended in a 5 ml volume of fixative master mix of Paraformaldehyde-Glutaraldehyde Solution Karnovsky’s Fixative (Electron Microscopy Sciences). Fixative solution was prepared with 16% paraformaldehyde solution, 5% Glutaraldehyde EM Grade, and 0.2 M Sodium phosphate buffer. Fixed samples were outsourced to Charles River laboratories for EM imaging service (Frederick PAI Project No. 20354508). Each sample was examined on a transmission electron microscope (TEM), and up to 10 representative digital images per sample were taken with an AMT camera.

### Immunofluorescence

Primary glioblastoma stem cells were seeded in complete media without heparin on cell-culture treated chamber slides (Ibidi) overnight and transfected after 24 hours with x2 lipid (Mirus Bio) and either 25 nM control (Horizon) or HDAC7 (Horizon) siRNA. All immunofluorescence staining was performed 48 hours post-transfection.

For H3.3, cells were fixed/permeabilized with 100% ice-cold methanol for 5 min and incubated in blocking buffer (1% BSA, 10% donkey serum, 22.52 mg/mL glycine in PBST (PBS+ 0.1% Tween 20)) for 1 hour at room temperature. Primary antibody (Abcam) was applied 1:1000 in primary antibody buffer (1% BSA in PBST) overnight at 4°C. Cells were washed with DPBS and incubated with secondary antibody (Jackson) 1:100 in DPBS for 1 hour and mounted with DAPI (Vector Labs).

For H3k9me3, cells were fixed with 4% paraformaldehyde (methanol-free) for 10 min, permeabilized with .1% Triton in DPBS for 5 min. Blocking, primary antibody (Active Motif), secondary antibody, and mounting was applied as described above.

Images were taken using a Zeiss Axiovert Inverted Microscope with Apotome II.

### H3K9me3 CHIP-seq

GSCs were seeded, transfected with siRNA against HDAC7 for 48 hours then processed for Chip-Seq using the standard active motif ChIP-IT ChIP-Seq protocol. Cells reached a confluency of 80% post transfection and were then fixed in 1% formaldehyde. Following fixation, samples were sonicated, and sheared chromatin was immunoprecipitated using a ChIP validated H3K9me3 antibody (Active Motif). The ChIP DNA was then reversed cross-linked, and DNA was collected for library preparation and sequencing.

### ChIP-Seq Analysis

Following library preparation, the 75-nt single-end (SE75) sequence reads generated by Illumina sequencing were mapped to the genome using the BWA algorithm with default settings and filtering out reads with more than 2 mismatches and with no unique alignment. The reads were extended to a length of 200 bp using the active motif software then subsequently divided into bins and stored as a bigwig file. Following read alignment and density identification, SICER was used for peak calling at default settings (FDR 1e-10 with gap parameter of 600 bp). Peak filtering was performed by removing false ChIP-Seq peaks within the ENCODE blacklist. Spike-in chromatin was performed, and number of test tags adjusted for normalization. For annotation, peaks were associated to genes if found within 10kb upstream/downstream limit.

### Proximity Ligation Assay

Cells were seeded at a density of 2.5 * 10^4 cells on fibronectin-coated 10mm German Glass coverslips placed in 4-well plates and cultured for 24 hours. The cells were washed once with PBS, fixed in methanol for 5 minutes at room temperature, then blocked for an hour using the Duolink Blocking Solution. All subsequent washes were in quantities of 4 mL at room temperature, with the coverslips transferred to 6 well plates. All reactions and incubations occurred on parafilm taped inside a humidity chamber.

The HDAC7 antibody was added in a 1:100 dilution into Duolink Antibody Diluent, 50 µL was placed on each coverslip, and incubated overnight at 4°C. The coverslips were washed 3 times in 4mL 1x Wash Buffer A for 5 minutes each. The anti-rabbit PLUS probe was diluted 1:5 in the Duolink Antibody Diluent, 40 µL was applied to each coverslip, and incubated for 1 hour at 37°C. Coverslips were washed 3 times with 1x Wash Buffer A for 5 minutes. The conjugated H3.3-MINUS antibody (created with the Duolink Probemaker MINUS) was diluted 1:300 in PLA probe diluent, 50 µL was added to each coverslip, and incubated overnight at 4°C. Coverslips were washed 3 times with 1x Wash Buffer A for 5 minutes. The 5x Duolink ligation buffer was diluted 1:5 in high purity water, and the Ligase was added in a 1:40 dilution right before adding 40 µL of the mix to the coverslips. These were incubated for 30 minutes at 37°C. The coverslips were washed 3 times with 1x Wash Buffer A for 5 minutes. The 5x Duolink amplification buffer was diluted 1:5 in high purity water, and the Polymerase was added in a 1:80 dilution right before applying 40 µL of the amplification solution to the coverslips, and incubated for 100 minutes at 37°C. The amplification buffer is light-sensitive, and was protected from the light throughout, as well as for all subsequent steps. The slides were washed 2 times in 1x Wash Buffer B for 10 minutes, then 1 time in 0.01x Wash Buffer B for 1 minute. Two coverslips were mounted onto each slide, using the Vectashield Mounting Medium with DAPI. They were left to dry for at least one hour, and imaged using the 83x objective on the Zeiss Axio Imager with the ApoTome 2.0. Slides were stored at -20°C. Negative controls had either the conjugated H3.3-MINUS antibody or the HDAC7 antibody excluded.

### DNA Fiber Assay

GSCs were transfected with siControl or siHDAC7, as above. DNA Fiber Assays were carried out according to published protocols^59^. Quantification of fibers from three images obtained from a total of two independent experiments was completed using semi-automated technique available via the DNA Stranding software package^60^. In each group, individual fibers were segmented automatically using the software’s pre-specified input parameters and selecting the 20-minute pulse labelling of CIdU and IdU according to the DNA fiber assay protocol. After automatic segmentation, manual review of individual fiber labels was conducted to remove erroneously selected fiber segmentations and correct apparent mistakes, in line with the published methods.

N=291 fibers from cells transfected with siControl and n=167 fibers from cells transfected with siHDAC7 were identified and quantified in the final analysis. Quantitative comparison of replication fork speed and CIdU / IdU ratio metrics was carried out in GraphPad Prism.

### Cell Cycle Analysis using Flow Cytometry

Cells were transfected as above. 48 hours after transfection, 300,000 cells of each sample were passaged onto a new 6-well plate and allowed to grow for 48 hours. Media was aspirated from each well, and washed with 500 µL PBS. Cells were lifted using 500 µL of StemPro™ Accutase™, and combined the two associated wells into one low adhesion tube. Wells were washed with 500 µL PBS and collected into those same tubes. All centrifugations were performed with a swinging bucket rotor. The tubes were centrifuged at 230 g for 5 minutes. After aspirating all but the cell pellet, the cells were fixed with 1 mL ice cold 70% ethanol for 30 minutes, on ice. The ethanol was added drop wise to the pellet while vortexing to minimize clumping. Cells were washed 3 times in PBS with 0.5% BSA to prevent cells sticking to the cell walls, followed by centrifugation at 850 g for 5 minutes. Cells were counted prior to the final spin, placing 500,000 cells into each tube. Cells were resuspended in 250 µL of FxCycle PI/RNase Staining Solution and analyzed using flow cytometry.

### Chromatin lysate isolation for **γ**H2AX detection

First, CSK buffer was prepared with a final pH=6.8 and consisting of the following components, in water: 10mM PIPES, 100mM NaCl, 300mM Sucrose, 3mM MgCl2, 1mM EGTA, and 0.2% Triton X-100. Buffer was stored at 4°C until ready for isolation. After plating GSCs on 6-well clear-bottomed plates in transfection media (350k / well), cells were transfected after 24h with siCTRL or siHDAC7. Cell lysates were then collected after 48 hours. Additional siHDAC7-transfected cells were treated with etoposide (50 µM) for collection 30 mins and 120 mins after treatment. For collection of lysates, cells from individual wells were detached from the plate using StemPro Accutase, spun for 5 minutes at 230g to pellet, and the pellet was resuspended in 500 µL of CSK Buffer on ice for 30 minutes, vortexing samples every 5-10 minutes. Samples were then centrifuged at 4°C for 10 minutes at 10,000 g. After aspirating the supernatant containing chromatin-unbound proteins and leaving only chromatin-bound protein as a pellet, each pellet was washed twice again with CSK buffer. The pellet was then re-suspended in 100µL Laemmli buffer (99µL 1% SDS plus 1µL phosphatase inhibitor cocktail). Samples were then sonicated and heated to 95°C to denature proteins in preparation for Western Blotting.

## QUANTIFICATION AND STATISTICAL ANALYSIS

To determine significance among the means of two independent groups, we perform an unpaired two-tailed t test. To verify Gaussian distribution of data before applying the t test, we perform the D’Agostino and Pearson and Shapiro-Wilk normality tests.

To calculate sample size, we assume the following:

**Input:** Tails= two, Effect size Delta= 2.5, α error probe= 0.05, Power (1-β err probe)= 0.90, Allocation ration N2/N1= 1

**Output:** Noncentrality parameter δ= 3.9528471, Critical t= 2.3, Df=8

### Actual Power= 0.9

Significance is stated as follows: (ns), p < 0.05 (*), p < 0.01 (**), p < 0.001 (***), p < 0.0001 (****).

## RESOURCE AVAILABILITY

NGS and Nanostring data included in this study are submitted to GEO: https://www.ncbi.nlm.nih.gov/geo/query/acc.cgi?acc=GSE 286178 Token for GEO review: **ezcnumeynjwphyl**

Further information and requests for resources and reagents should be directed to and will be fulfilled by the lead contact, Nikos Tapinos.

## Author Contributions

O.H., M.P., O.L., J.P.Z., A.C., L.J.W., S.S., D.L., L.H., L.T., E.F., A.F., D.K.:

Experimentation, Data Analysis, Figures, Manuscript Writing and Editing

N.T.: Conceptualization, Supervision, Data Analysis, Writing of Manuscript, Funding

## Acknowledgments

The authors would like to thank the Department of Pathology at Rhode Island Hospital for providing the 84 glioblastoma tissue samples for Nanostring Analysis. We also thank Charles River for Electron Microscopy services. This study was supported by a Warren Alpert Foundation Grant #17775 to N.T., private philanthropic donations to the Laboratory of Cancer Epigenetics and Plasticity and from internal support of Brown University to N.T.

## References

1. Wainwright, E.N., and Scaffidi, P. (2017). Epigenetics and Cancer Stem Cells: Unleashing, Hijacking, and Restricting Cellular Plasticity. Trends Cancer 3, 372–386. 10.1016/j.trecan.2017.04.004.

2. Guetta-Terrier, C., Karambizi, D., Akosman, B., Zepecki, J.P., Chen, J.S., Kamle, S., Fajardo, J.E., Fiser, A., Singh, R., Toms, S.A., et al. (2023). Chi3l1 Is a Modulator of Glioma Stem Cell States and a Therapeutic Target in Glioblastoma. Cancer Res 83, 1984–1999. 10.1158/0008-5472.CAN-21-3629.

3. Allfrey, V.G., Faulkner, R., and Mirsky, A.E. (1964). Acetylation and Methylation of Histones and Their Possible Role in the Regulation of Rna Synthesis. Proc Natl Acad Sci U S A 51, 786–794. 10.1073/pnas.51.5.786.

4. Li, G., Tian, Y., and Zhu, W.G. (2020). The Roles of Histone Deacetylases and Their Inhibitors in Cancer Therapy. Front Cell Dev Biol 8, 576946. 10.3389/fcell.2020.576946.

5. Park, S.Y., and Kim, J.S. (2020). A short guide to histone deacetylases including recent progress on class II enzymes. Exp Mol Med 52, 204–212. 10.1038/s12276-020-0382-4.

6. Falkenberg, K.J., and Johnstone, R.W. (2014). Histone deacetylases and their inhibitors in cancer, neurological diseases and immune disorders. Nat Rev Drug Discov 13, 673–691. 10.1038/nrd4360.

7. West, A.C., and Johnstone, R.W. (2014). New and emerging HDAC inhibitors for cancer treatment. J Clin Invest 124, 30–39. 10.1172/JCI69738.

8. Stypula-Cyrus, Y., Damania, D., Kunte, D.P., Cruz, M.D., Subramanian, H., Roy, H.K., and Backman, V. (2013). HDAC up-regulation in early colon field carcinogenesis is involved in cell tumorigenicity through regulation of chromatin structure. PLoS One 8, e64600. 10.1371/journal.pone.0064600.

9. Gao, S., Liu, H., Hou, S., Wu, L., Yang, Z., Shen, J., Zhou, L., Zheng, S.S., and Jiang, B. (2018). MiR-489 suppresses tumor growth and invasion by targeting HDAC7 in colorectal cancer. Clin Transl Oncol 20, 703–712. 10.1007/s12094-017-1770-7.

10. Ouaissi, M., Sielezneff, I., Silvestre, R., Sastre, B., Bernard, J.P., Lafontaine, J.S., Payan, M.J., Dahan, L., Pirro, N., Seitz, J.F., et al. (2008). High histone deacetylase 7 (HDAC7) expression is significantly associated with adenocarcinomas of the pancreas. Ann Surg Oncol 15, 2318–2328. 10.1245/s10434-008-9940-z.

11. 11. Ouaissi, M., Silvy, F., Loncle, C., Ferraz da Silva, D., Martins Abreu, C., Martinez, E., Berthezene, P., Cadra, S., Le Treut, Y.P., Hardwigsen, J., et al. (2014). Further characterization of HDAC and SIRT gene expression patterns in pancreatic cancer and their relation to disease outcome. PLoS One 9, e108520. 10.1371/journal.pone.0108520.

12. Freese, K., Seitz, T., Dietrich, P., Lee, S.M.L., Thasler, W.E., Bosserhoff, A., and Hellerbrand, C. (2019). Histone Deacetylase Expressions in Hepatocellular Carcinoma and Functional Effects of Histone Deacetylase Inhibitors on Liver Cancer Cells In Vitro. Cancers (Basel) 11. 10.3390/cancers11101587.

13. Yu, Y., Cao, F., Yu, X., Zhou, P., Di, Q., Lei, J., Tai, Y., Wu, H., Li, X., Wang, X., et al. (2017). The expression of HDAC7 in cancerous gastric tissues is positively associated with distant metastasis and poor patient prognosis. Clin Transl Oncol 19, 1045–1054. 10.1007/s12094-017-1639-9.

14. Wei, Y., Zhou, F., Zhou, H., Huang, J., Yu, D., and Wu, G. (2018). Endothelial progenitor cells contribute to neovascularization of non-small cell lung cancer via histone deacetylase 7-mediated cytoskeleton regulation and angiogenic genes transcription. Int J Cancer 143, 657–667. 10.1002/ijc.31349.

15. Wu, M.Y., Fu, J., Xiao, X., Wu, J., and Wu, R.C. (2014). MiR-34a regulates therapy resistance by targeting HDAC1 and HDAC7 in breast cancer. Cancer Lett 354, 311–319. 10.1016/j.canlet.2014.08.031.

16. Yu, X., Wang, M., Wu, J., Han, Q., and Zhang, X. (2019). ZNF326 promotes malignant phenotype of glioma by up-regulating HDAC7 expression and activating Wnt pathway. J Exp Clin Cancer Res 38, 40. 10.1186/s13046-019-1031-4.

17. Peixoto, P., Blomme, A., Costanza, B., Ronca, R., Rezzola, S., Palacios, A.P., Schoysman, L., Boutry, S., Goffart, N., Peulen, O., et al. (2016). HDAC7 inhibition resets STAT3 tumorigenic activity in human glioblastoma independently of EGFR and PTEN: new opportunities for selected targeted therapies. Oncogene 35, 4481–4494. 10.1038/onc.2015.506.

18. Ahmad, K., and Henikoff, S. (2002). The histone variant H3.3 marks active chromatin by replication-independent nucleosome assembly. Mol Cell 9, 1191–1200. 10.1016/s1097-2765(02)00542-7.

19. Schwartz, B.E., and Ahmad, K. (2005). Transcriptional activation triggers deposition and removal of the histone variant H3.3. Genes Dev 19, 804–814. 10.1101/gad.1259805.

20. Szenker, E., Ray-Gallet, D., and Almouzni, G. (2011). The double face of the histone variant H3.3. Cell Res 21, 421–434. 10.1038/cr.2011.14.

21. Deal, R.B., Henikoff, J.G., and Henikoff, S. (2010). Genome-wide kinetics of nucleosome turnover determined by metabolic labeling of histones. Science 328, 1161–1164. 10.1126/science.1186777.

22. MacAlpine, H.K., Gordan, R., Powell, S.K., Hartemink, A.J., and MacAlpine, D.M. (2010). Drosophila ORC localizes to open chromatin and marks sites of cohesin complex loading. Genome Res 20, 201–211. 10.1101/gr.097873.109.

23. Eaton, M.L., Prinz, J.A., MacAlpine, H.K., Tretyakov, G., Kharchenko, P.V., and MacAlpine, D.M. (2011). Chromatin signatures of the Drosophila replication program. Genome Res 21, 164–174. 10.1101/gr.116038.110.

24. Stroud, H., Otero, S., Desvoyes, B., Ramirez-Parra, E., Jacobsen, S.E., and Gutierrez, C. (2012). Genome-wide analysis of histone H3.1 and H3.3 variants in Arabidopsis thaliana. Proc Natl Acad Sci U S A 109, 5370–5375. 10.1073/pnas.1203145109.

25. Paranjape, N.P., and Calvi, B.R. (2016). The Histone Variant H3.3 Is Enriched at Drosophila Amplicon Origins but Does Not Mark Them for Activation. G3 (Bethesda) 6, 1661–1671. 10.1534/g3.116.028068.

26. Strobino, M., Wenda, J.M., Padayachy, L., and Steiner, F.A. (2020). Loss of histone H3.3 results in DNA replication defects and altered origin dynamics in C. elegans. Genome Res 30, 1740–1751. 10.1101/gr.260794.120.

27. Adam, S., Polo, S.E., and Almouzni, G. (2013). Transcription recovery after DNA damage requires chromatin priming by the H3.3 histone chaperone HIRA. Cell 155, 94–106. 10.1016/j.cell.2013.08.029.

28. Clement, C., Orsi, G.A., Gatto, A., Boyarchuk, E., Forest, A., Hajj, B., Mine-Hattab, J., Garnier, M., Gurard-Levin, Z.A., Quivy, J.P., and Almouzni, G. (2018). High-resolution visualization of H3 variants during replication reveals their controlled recycling. Nat Commun 9, 3181. 10.1038/s41467-018-05697-1.

29. Shi, L., Wen, H., and Shi, X. (2017). The Histone Variant H3.3 in Transcriptional Regulation and Human Disease. J Mol Biol 429, 1934–1945. 10.1016/j.jmb.2016.11.019.

30. Goldberg, A.D., Banaszynski, L.A., Noh, K.M., Lewis, P.W., Elsaesser, S.J., Stadler, S., Dewell, S., Law, M., Guo, X., Li, X., et al. (2010). Distinct factors control histone variant H3.3 localization at specific genomic regions. Cell 140, 678–691. 10.1016/j.cell.2010.01.003.

31. Drane, P., Ouararhni, K., Depaux, A., Shuaib, M., and Hamiche, A. (2010). The death-associated protein DAXX is a novel histone chaperone involved in the replication-independent deposition of H3.3. Genes Dev 24, 1253–1265. 10.1101/gad.566910.

32. Lewis, P.W., Elsaesser, S.J., Noh, K.M., Stadler, S.C., and Allis, C.D. (2010). Daxx is an H3.3- specific histone chaperone and cooperates with ATRX in replication-independent chromatin assembly at telomeres. Proc Natl Acad Sci U S A 107, 14075–14080. 10.1073/pnas.1008850107.

33. Wang, S., Fairall, L., Pham, T.K., Ragan, T.J., Vashi, D., Collins, M.O., Dominguez, C., and Schwabe, J.W.R. (2023). A potential histone-chaperone activity for the MIER1 histone deacetylase complex. Nucleic Acids Res 51, 6006–6019. 10.1093/nar/gkad294.

34. Martin, M., Kettmann, R., and Dequiedt, F. (2007). Class IIa histone deacetylases: regulating the regulators. Oncogene 26, 5450–5467. 10.1038/sj.onc.1210613.

35. Kao, H.Y., Verdel, A., Tsai, C.C., Simon, C., Juguilon, H., and Khochbin, S. (2001). Mechanism for nucleocytoplasmic shuttling of histone deacetylase 7. J Biol Chem 276, 47496–47507. 10.1074/jbc.M107631200.

36. Loyola, A., Tagami, H., Bonaldi, T., Roche, D., Quivy, J.P., Imhof, A., Nakatani, Y., Dent, S.Y., and Almouzni, G. (2009). The HP1alpha-CAF1-SetDB1-containing complex provides H3K9me1 for Suv39-mediated K9me3 in pericentric heterochromatin. EMBO Rep 10, 769–775. 10.1038/embor.2009.90.

37. Dong, J., LeBlanc, C., Poulet, A., Mermaz, B., Villarino, G., Webb, K.M., Joly, V., Mendez, J., Voigt, P., and Jacob, Y. (2021). H3.1K27me1 maintains transcriptional silencing and genome stability by preventing GCN5-mediated histone acetylation. Plant Cell 33, 961–979. 10.1093/plcell/koaa027.

38. Ferrari, K.J., Scelfo, A., Jammula, S., Cuomo, A., Barozzi, I., Stutzer, A., Fischle, W., Bonaldi, T., and Pasini, D. (2014). Polycomb-dependent H3K27me1 and H3K27me2 regulate active transcription and enhancer fidelity. Mol Cell 53, 49–62. 10.1016/j.molcel.2013.10.030.

39. 39. Warnon, C., Bouhjar, K., Ninane, N., Verhoyen, M., Fattaccioli, A., Fransolet, M., Lambert de Rouvroit, C., Poumay, Y., Piel, G., Mottet, D., and Debacq-Chainiaux, F. (2021). HDAC2 and 7 down-regulation induces senescence in dermal fibroblasts. Aging (Albany NY) 13, 17978–18005. 10.18632/aging.203304.

40. Zhu, C., Chen, Q., Xie, Z., Ai, J., Tong, L., Ding, J., and Geng, M. (2011). The role of histone deacetylase 7 (HDAC7) in cancer cell proliferation: regulation on c-Myc. Journal of molecular medicine 89, 279–289. 10.1007/s00109-010-0701-7.

41. Maya-Mendoza, A., Moudry, P., Merchut-Maya, J.M., Lee, M., Strauss, R., and Bartek, J. (2018). High speed of fork progression induces DNA replication stress and genomic instability. Nature 559, 279–284. 10.1038/s41586-018-0261-5.

42. Lahm, A., Paolini, C., Pallaoro, M., Nardi, M.C., Jones, P., Neddermann, P., Sambucini, S., Bottomley, M.J., Lo Surdo, P., Carfi, A., et al. (2007). Unraveling the hidden catalytic activity of vertebrate class IIa histone deacetylases. Proc Natl Acad Sci U S A 104, 17335–17340. 10.1073/pnas.0706487104.

43. Brancolini, C., Di Giorgio, E., Formisano, L., and Gagliano, T. (2021). Quis Custodiet Ipsos Custodes (Who Controls the Controllers)? Two Decades of Studies on HDAC 9. Life (Basel) 11. 10.3390/life11020090.

44. 44. Di Giorgio, E., and Brancolini, C. (2016). Regulation of class IIa HDAC activities: it is not only matter of subcellular localization. Epigenomics 8, 251–269. 10.2217/epi.15.106.

45. Brancolini, C., Gagliano, T., and Minisini, M. (2022). HDACs and the epigenetic plasticity of cancer cells: Target the complexity. Pharmacol Ther 238, 108190. 10.1016/j.pharmthera.2022.108190.

46. Witt, A.E., Lee, C.W., Lee, T.I., Azzam, D.J., Wang, B., Caslini, C., Petrocca, F., Grosso, J., Jones, M., Cohick, E.B., et al. (2017). Identification of a cancer stem cell-specific function for the histone deacetylases, HDAC1 and HDAC7, in breast and ovarian cancer. Oncogene 36, 1707-1720. 10.1038/onc.2016.337.

47. Caslini, C., Hong, S., Ban, Y.J., Chen, X.S., and Ince, T.A. (2019). HDAC7 regulates histone 3 lysine 27 acetylation and transcriptional activity at super-enhancer-associated genes in breast cancer stem cells. Oncogene 38, 6599–6614. 10.1038/s41388-019-0897-0.

48. Tagami, H., Ray-Gallet, D., Almouzni, G., and Nakatani, Y. (2004). Histone H3.1 and H3.3 complexes mediate nucleosome assembly pathways dependent or independent of DNA synthesis. Cell 116, 51–61. 10.1016/s0092-8674(03)01064-x.

49. Hathaway, N.A., Bell, O., Hodges, C., Miller, E.L., Neel, D.S., and Crabtree, G.R. (2012). Dynamics and memory of heterochromatin in living cells. Cell 149, 1447–1460. 10.1016/j.cell.2012.03.052.

50. Kunkel, T.A. (2004). DNA replication fidelity. J Biol Chem 279, 16895–16898. 10.1074/jbc.R400006200.

51. Sedlackova, H., Rask, M.B., Gupta, R., Choudhary, C., Somyajit, K., and Lukas, J. (2020). Equilibrium between nascent and parental MCM proteins protects replicating genomes. Nature 587, 297–302. 10.1038/s41586-020-2842-3.

52. Zepecki, J.P., Snyder, K.M., Moreno, M.M., Fajardo, E., Fiser, A., Ness, J., Sarkar, A., Toms, S.A., and Tapinos, N. (2018). Regulation of human glioma cell migration, tumor growth, and stemness gene expression using a Lck targeted inhibitor. Oncogene. 10.1038/s41388-018-0546-z.

53. Menyhart, O., Nagy, A., and Gyorffy, B. (2018). Determining consistent prognostic biomarkers of overall survival and vascular invasion in hepatocellular carcinoma. R Soc Open Sci 5, 181006. 10.1098/rsos.181006.

54. Bowman, R.L., Wang, Q., Carro, A., Verhaak, R.G., and Squatrito, M. (2017). GlioVis data portal for visualization and analysis of brain tumor expression datasets. Neuro Oncol 19, 139–141. 10.1093/neuonc/now247.

55. Love, M.I., Huber, W., and Anders, S. (2014). Moderated estimation of fold change and dispersion for RNA-seq data with DESeq2. Genome Biol 15, 550. 10.1186/s13059-014- 0550-8.

56. Johnson, S., Chen, H., and Lo, P.K. (2013). In vitro Tumorsphere Formation Assays. Bio Protoc 3. 10.21769/bioprotoc.325.

57. 57. MacLean, B., Tomazela, D.M., Shulman, N., Chambers, M., Finney, G.L., Frewen, B., Kern, R., Tabb, D.L., Liebler, D.C., and MacCoss, M.J. (2010). Skyline: an open source document editor for creating and analyzing targeted proteomics experiments. Bioinformatics 26, 966–968. 10.1093/bioinformatics/btq054.

58. Mohammed, H., Taylor, C., Brown, G.D., Papachristou, E.K., Carroll, J.S., and D’Santos, C.S. (2016). Rapid immunoprecipitation mass spectrometry of endogenous proteins (RIME) for analysis of chromatin complexes. Nat Protoc 11, 316–326. 10.1038/nprot.2016.020.

59. Halliwell, J.A., Gravells, P., and Bryant, H.E. (2020). DNA Fiber Assay for the Analysis of DNA Replication Progression in Human Pluripotent Stem Cells. Curr Protoc Stem Cell Biol 54, e115. 10.1002/cpsc.115.

60. Li, L., Kolinjivadi, A.M., Ong, K.H., Young, D.M., Marini, G.P.L., Chan, S.H., Chong, S.T., Chew, E.L., Lu, H., Gole, L., et al. (2022). Automatic DNA replication tract measurement to assess replication and repair dynamics at the single-molecule level. Bioinformatics 38, 4395–4402. 10.1093/bioinformatics/btac533.

