## Supplemental Figure Legends for "Inhibition of HDAC7 reprograms the histone H3.3 landscape to induce heterochromatin spreading and DNA replication defects in cancer cells"

**Supplementary Figure Legends**

**Figure S1: Class specificity of HDAC7 targeting siRNAs.** A) Functional classification of the significantly increased genes shows that they have significant pro-tumorigenic roles like Vimentin, Lox, SERPINH1, TGFb. B & C) Four independent siRNAs targeting HDAC7 were designed, and class specificity was examined in GSCs. All siRNAs significantly inhibit expression of HDAC7 (***p<0.005, ****p<0.001). siHDAC7 siRNAs-1, 3 & 4 do not show class specificity since siHDAC-1 significantly increases expression of HDAC9 (*p<0.05), siHDAC7-3 and siHDAC7-4 significantly inhibit expression of HDAC5 (*p<0.05 and **p<0.01 respectively). siHDAC7-2 shows class specificity and significant inhibition of HDAC7 (data included in the main paper).

**Figure S2: Baseline Histone modifications in patients derived GSCs.**

At baseline (siControl GSCs), dimethylation of histone marks shows the highest abundance in comparison to other modifications in GSCs. The most modified histone sites were H3K9, H3.1K27, H3.3K27 and H4K20.

**Figure S3: Downregulation of cancer stemness genes and effect on cancer stem cell self-renewal of GSCs and effect on EdU incorporation in PANC-1 cells.** A) Plot showing significant downregulation of genes with central role in cancer stem cell biology following inhibition of HDAC7 expression. B) GSEA following inhibition of HDAC7 expression shows significant inhibition of embryonic stem cell signatures and FOXR2 target genes. C) Tumor sphere formation assay shows significant (n=3, p<0.0001) inhibition of GSC self- renewal following inhibition of HDAC7 expression. D) Inhibition of HDAC7 expression in the pancreatic cancer cell line PANC-1 results in significant inhibition of EdU incorporation (n=3, p<0.05).

**Figure S4: Inhibition of HDAC7 does not induce senescence phenotype and sensitizes cancer cells to Camptothecin.** A) Senescence Gene expression signature (GSEA) from RNA-seq of GSCs treated with siControl or siHDAC7 shows that inhibition of HDAC7 expression does not induce expression of senescent genes. B) Representative phase contrast images at 20X magnification of beta-galactosidase expression (blue) in GSCs treated with siControl or siHDAC7. C) Quantification of beta-galactosidase expression from 550 cells treated with siControl or siHDAC7. Results are presented as percent of positive cells relative to total number of cells. D) Addition of 0.25uM or 5uM Camptothecin to siHDAC7 treated cells significantly increases the growth inhibitory effect of Camptothecin alone (n=3 biological replicates, p<0.05).

**Supplementary Table 1: List of genes associated with genomic sites with increased H3K9me3 following inhibition of HDAC7.**
