## Supplementary figures and images for "Inhibition of HDAC7 reprograms the histone H3.3 landscape to induce heterochromatin spreading and DNA replication defects in cancer cells"

### Supplemental Figure 1

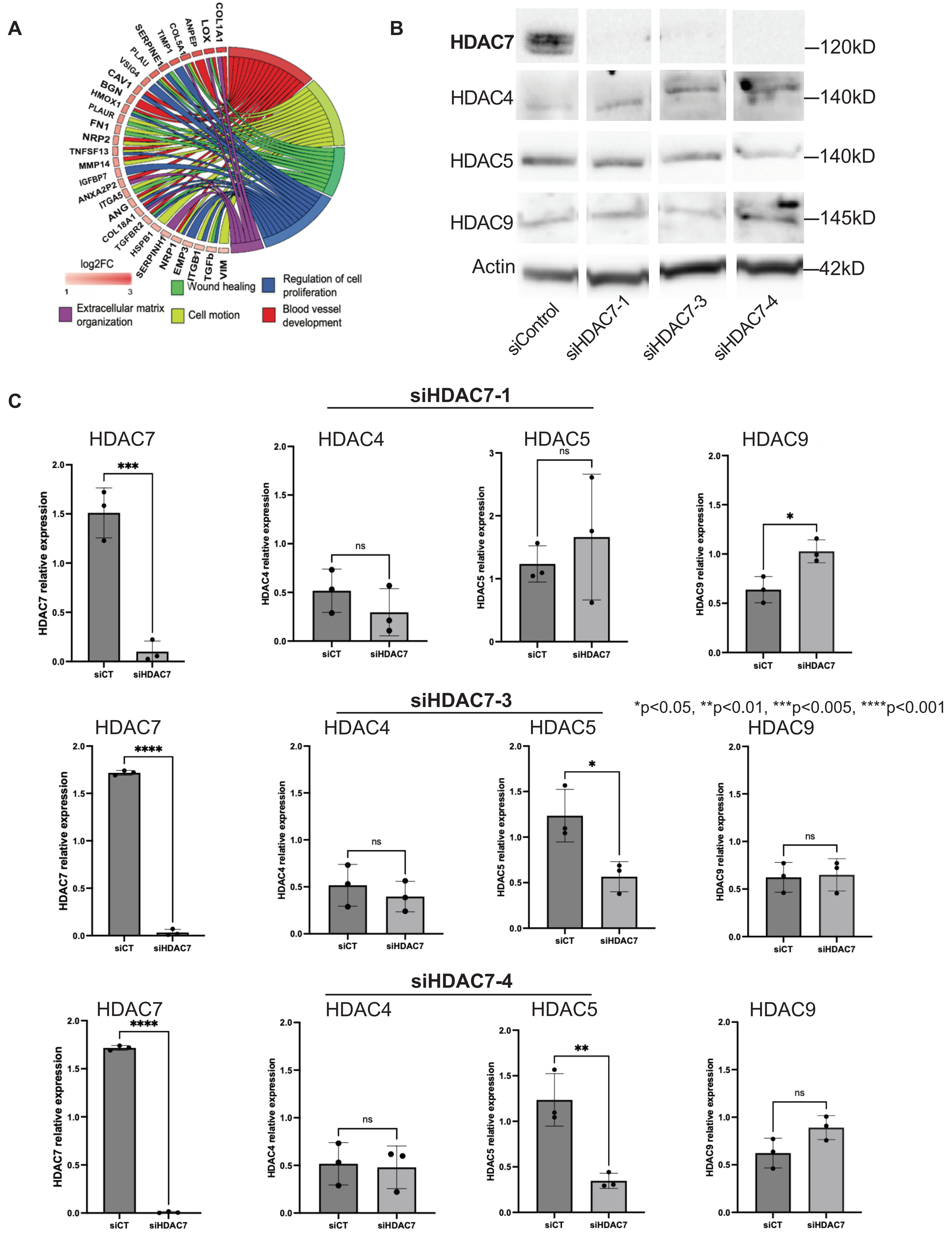

### Supplemental Figure 2

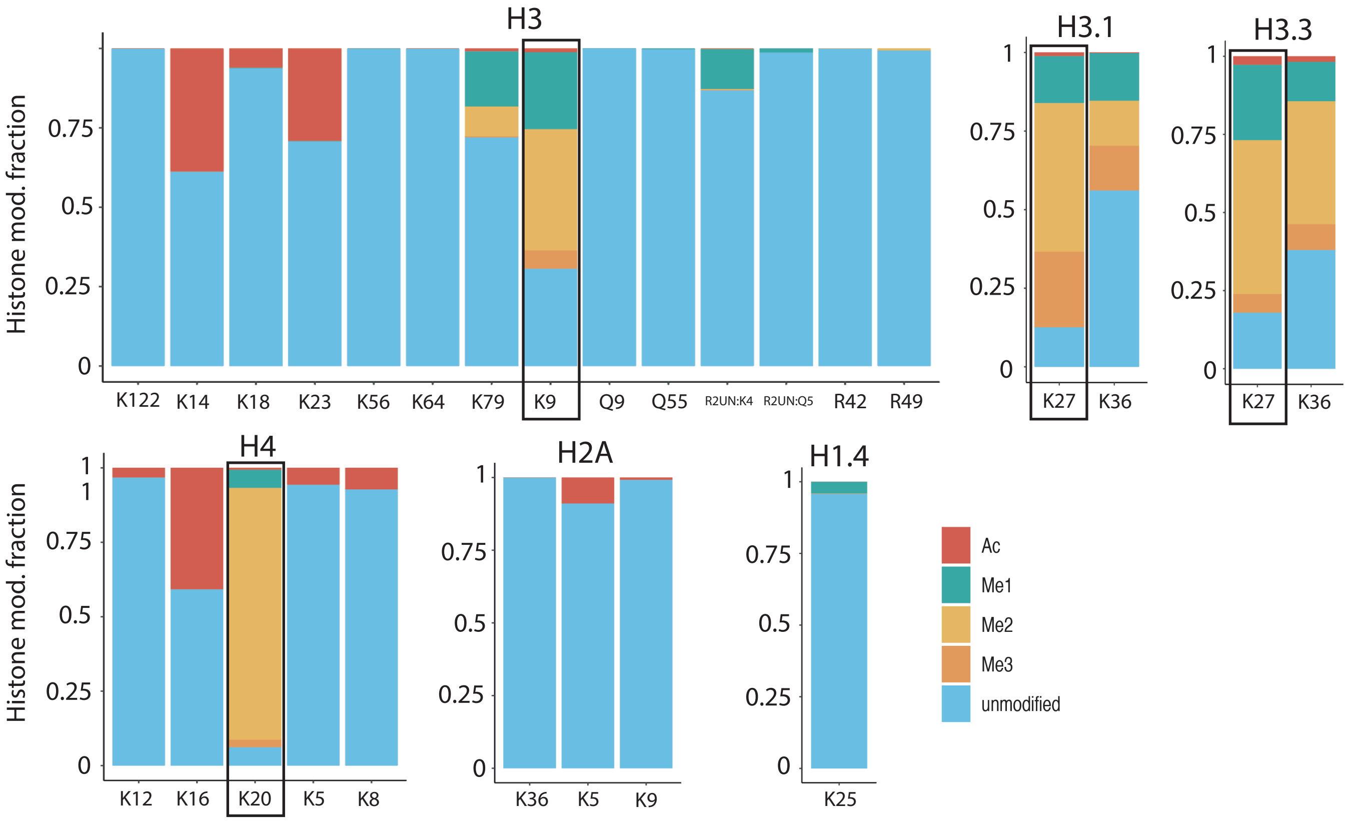

### Supplemental Figure 3

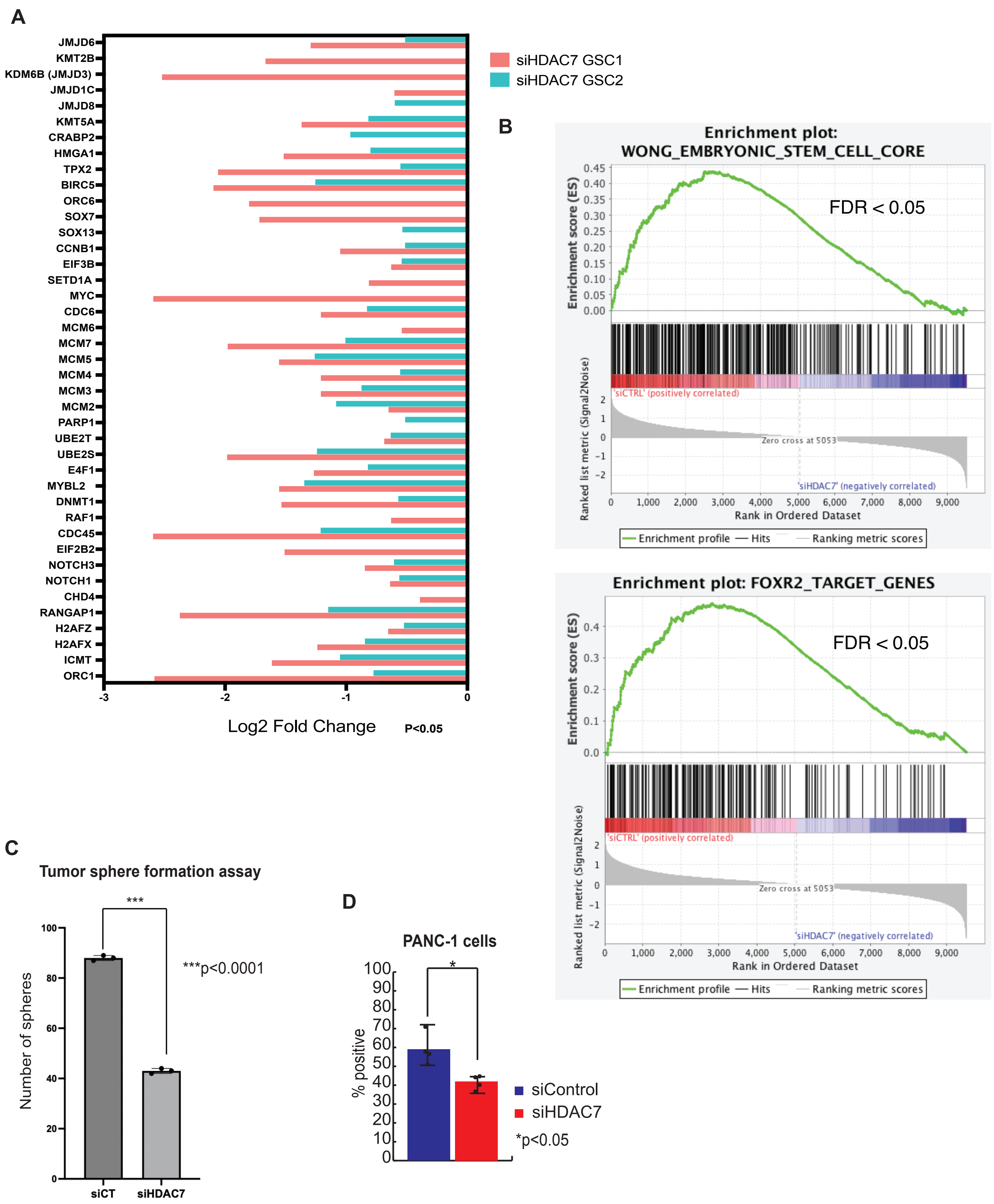

### Supplemental Figure 4

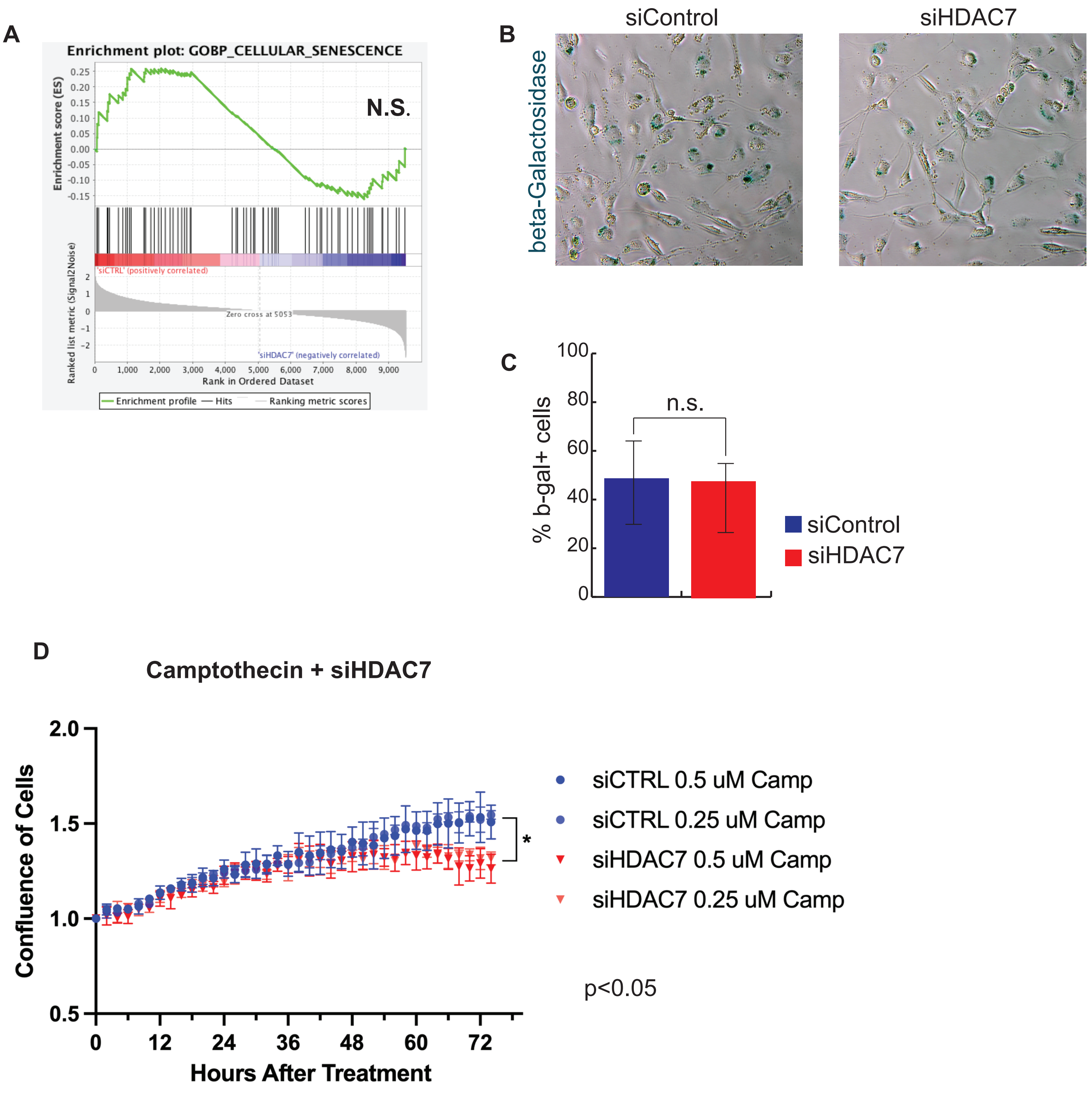
